# An Integrated muti-omics cell atlas of the human trabecular meshwork and ciliary body

**DOI:** 10.64898/2026.06.17.732980

**Authors:** Jinjing Jian, Xuan Bao, Jun Wang, Ye Zheng, Jin Li, Jianming Shao, Tingting Yang, Ismail Yaman, Jean Li, Han Chen, Nicholas Tolman, Richard H. Scheuermann, Jie J. Zheng, Carl Sheridan, Yutao Liu, Yiqin Du, Revathi Balasubramanian, Gulab S. Zode, Simon WM John, J Timothy Stout, W. Daniel Stamer, Yumei Li, Rui Chen

**Author notes:** Correspondence and requests for materials should be addressed to: Rui Chen,; Yumei Li,; W. Daniel Stamer,. Rui Chen is the lead contact. These authors contributed equally: Jinjing Jian, Xuan Bao, Jun Wang.

## Abstract

The trabecular meshwork (TM) and ciliary body (CB) regulate aqueous humor dynamics and intraocular pressure (IOP), and TM/Schlemm’s canal (SC) dysfunction underlies glaucoma. Here, we present a spatially resolved multi-omics atlas of human TM and CB, integrating snRNA-seq, scRNA-seq, and snATAC-seq from over one million cells and nuclei across 112 donors with Xenium spatial transcriptomics. We identified 9 major cell classes and 21 cell types, revealing heterogeneity, including undercharacterized fibroblast and epithelial subpopulations. Spatial mapping supported TM fibroblast zonation and CB epithelial organization. Regulatory analyses identified cell type–specific programs, including OTX/PAX networks in CB epithelium and SMAD3/TGF-β signaling in fibroblasts. Integration with glaucoma loci showed enrichment of non-coding variants in regulatory elements associated with POAG and PACG. Age- and ancestry-associated remodeling revealed divergent fibroblast aging with increased *PIEZO1*, suggesting impaired outflow and elevated IOP. Together, this high-resolution atlas links cellular, regulatory, and genetic variation to anterior segment function and glaucoma susceptibility.

## Introduction

Precise regulation of cellular composition and gene regulatory programs is fundamental to tissue function, yet remains incompletely understood in many human organs. Recent advances in single-cell and single-nucleus sequencing technologies have enabled the systematic dissection of cellular heterogeneity at unprecedented resolution, driving the construction of reference atlases across diverse tissues. These efforts, including those led by the Human Cell Atlas initiative, are transforming our understanding of human biology by linking cell identity to function and disease susceptibility^1, 2^.

In the human eye, large-scale transcriptomic studies have begun to map cellular diversity across both anterior and posterior segments^3, 4^, as well as the retina^5^. These atlases have revealed complex cellular ecosystems and highlighted both conserved and tissue-specific features across ocular structures. More recently, spatial transcriptomic approaches such as Xenium have further enabled the characterization of cellular organization and spatial architecture within ocular tissues^6^. However, despite these advances, key components of the anterior segment that are central to ocular physiology and disease remain insufficiently characterized, particularly at the level of cross-donor variation, spatial organization, and gene regulatory architecture.

The trabecular meshwork (TM) and ciliary body (CB) form a functionally coupled system that maintains intraocular pressure (IOP) by balancing aqueous humor outflow and production. The conventional outflow pathway, including the TM and Schlemm’s canal (SC) regulates fluid drainage through a specialized outflow pathway composed of heterogeneous cell populations with hybrid cellular, structural and contractile properties^7–9^. Dysfunction of conventional outflow homeostasis leads to elevated IOP and is a major contributor to glaucoma, a leading cause of irreversible blindness worldwide^10–12^. In contrast, the CB secretes aqueous humor into the eye through coordinated activity of its unique epithelial bilayer, which is the main target for glaucoma therapy^10, 13–15^. Although prior studies have identified major cell types within these tissues^3, 4, 12, 15, 16^, a comprehensive understanding of their cellular diversity, spatial organization, inter-tissue relationships, and regulatory programs remains lacking.

A major limitation of existing studies is the absence of integrated, multimodal, and cross-donor analyses that can simultaneously resolve transcriptional states and their underlying regulatory mechanisms. In particular, how chromatin accessibility landscapes shape cell identity in the TM and CB, how these features vary across individuals, how distinct cellular populations are spatially organized within anterior segment tissues, and how they relate to genetic risk for ocular disease remain largely unexplored. Addressing these gaps requires large-scale datasets and robust integration across modalities and donors.

Here, we present a comprehensive multi-omics atlas of the human trabecular meshwork and ciliary body (HTCCA), integrating 1,098,521 single nuclei from snRNA-seq, 332,995 single cells from scRNA-seq, spatial transcriptomic profiling using Xenium, and 628,567 nuclei profiled by snATAC-seq across a diverse cohort of donors. This integrated dataset resolves a broad spectrum of stromal, vascular, neural-associated, and immune cell populations, including fibroblasts, endothelial cells, pericytes, ciliary muscle cells, Schwann cells, melanocytes, and immune cells, consistent with prior observations in anterior segment tissues^4^. By combining transcriptomic, spatial, and chromatin accessibility data, we define both shared and tissue-enriched cellular states, characterize their spatial localization within TM and CB tissues, and uncover regulatory programs that distinguish the two structures.

Furthermore, this atlas enables systematic mapping of cell type–specific regulatory elements and transcription factors (TFs) networks, and provides a framework for linking genetic variation to cellular function through integration with GWAS and eQTL data. Together, our study establishes a comprehensive, high-resolution, multimodal reference for human TM and CB, offering new insights into the cellular, spatial, and regulatory basis of ocular physiology and disease.

## Results

### Single-cell and single-nucleus transcriptomic atlas of the human trabecular meshwork and ciliary body

To construct a comprehensive multi-omics atlas of the human TM (including the inner wall of SC) and CB, we performed single-nucleus RNA sequencing (snRNA-seq) and single-cell RNA sequencing (scRNA-seq) of isolated TM and CB tissue separately, following the workflow outlined in Fig. 1a. We integrated publicly available datasets^3, 4, 17^ with our newly generated data to generate high-resolution, well-curated human TM and CB atlas (Fig.1a, detailed in Methods). After quality control, a total of 1,098,521 nuclei from snRNA-seq and 332,995 cells from scRNA-seq across 112 healthy donors were retained (Fig. 1a; Supplementary Table 1). To capture variation across human populations, donors from diverse ancestry backgrounds were included, comprising 46% European, 27% African, 19% Hispanic, and 8% Asian individuals. Among these donors, 66% were male and 34% were female (Fig. 1b). Donors spanned a wide age range, from 0 to over 90 years (Fig. 1b). Given the known differences in gene expression profiles between scRNA-seq and snRNA-seq datasets, we constructed two separate atlases based on each modality. The snRNA-seq dataset was integrated with one published^4^, whereas the scRNA-seq dataset was integrated separately with two published datasets^3, 17^.

**Fig. 1.**
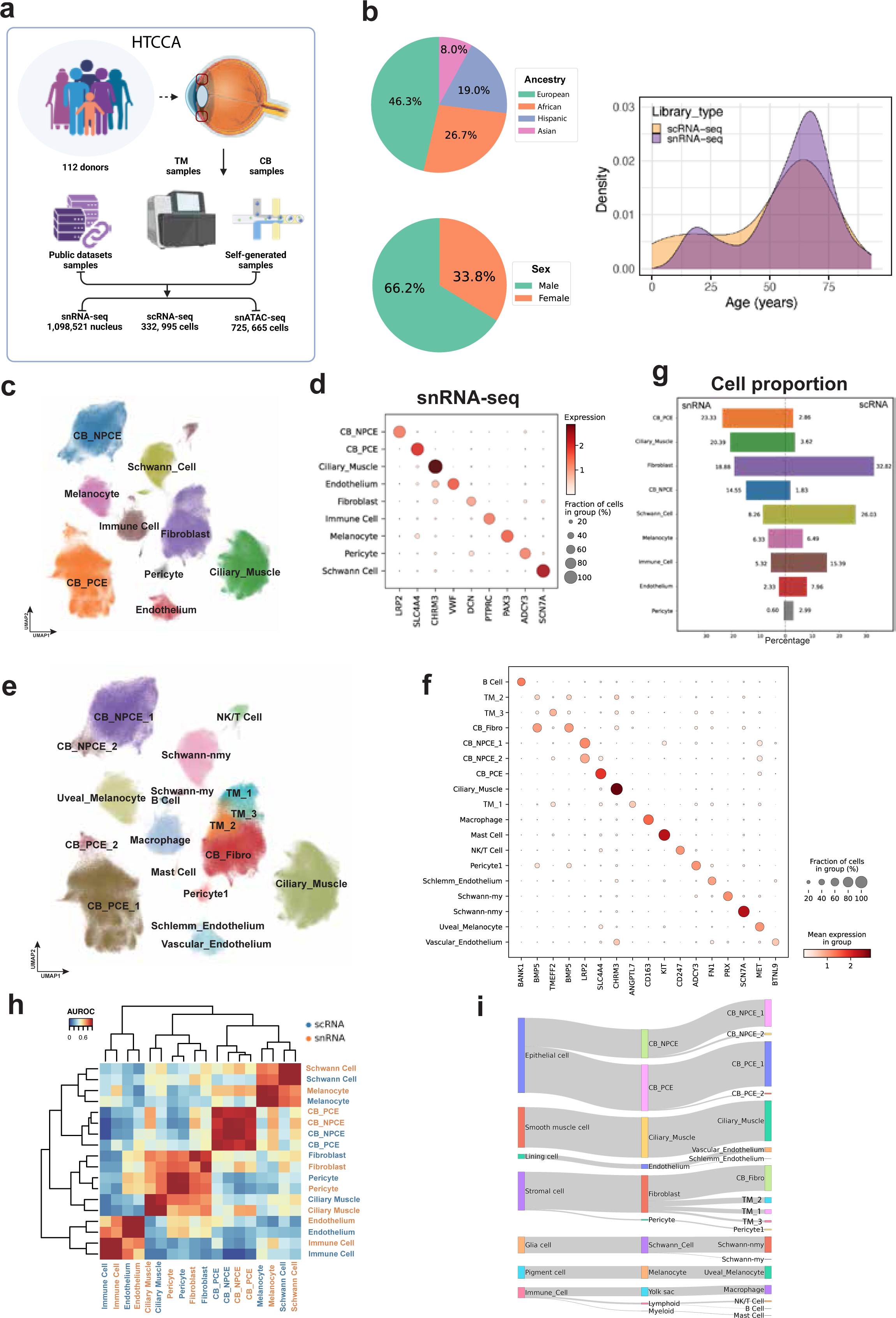
Overview of multi-omics atlas of the human TM and CB. **a,** Overview of the study design and dataset composition. Human TM and CB samples were collected from 112 donors and integrated with publicly available datasets. Both self-generated and public samples were profiled using snRNA-seq, scRNA-seq, and snATAC-seq, resulting in a large-scale multi-omic dataset comprising 110 snRNA-seq samples, 52 scRNA-seq samples, and 93 snATAC-seq samples. **b,** Demographic characteristics of donors included in this study. Pie charts show the proportion of cells contributed by different donor ethnicity and sex. Density plots depict the age distribution of donors across library types. **c,** The atlas of snRNA-seq datasets is visualized in a UMAP plot at a major class resolution, with cells colored based on their major classes. **d,** Comparison of major classes composition between snRNA-seq (left) and scRNA-seq (right) datasets. **e,** The atlas of scRNA-seq datasets is visualized in a UMAP plot at a major class resolution, with cells colored based on their major classes. **f,** Dot plots illustrating the distribution of expression levels of marker genes for major cell classes in snRNA-seq (left) and scRNA-seq (right) data. **g,** Cell type similarities of major classes between snRNA-seq (in orange) and scRNA-seq (in blue). The color key is the average AUROC of self-projection for cell classes. **h,** Sankey diagram illustrating the relationships among major cell groups (left column), cell classes (middle column), and cell type (right column).

Unsupervised clustering of the snRNA-seq data was performed and optimized using our established pipeline as described in the Methods^5^. In total, nine major cell clusters were identified and annotated based on known marker genes (Fig. 1d), including fibroblasts, endothelial cells, pericytes, ciliary muscle, pigmented and non-pigmented ciliary epithelium (PCE and NPCE), Schwann cells, melanocytes, and immune cells (Fig. 1c). The resulting cell annotations were consistent with those reported in previous studies^4, 17^ (Supplementary Table 1). Among these cells, 204,922 were derived from TM samples (46 donors), whereas 893,599 were from CB samples (82 donors). Several cell clusters, including immune cells, Schwann cells, melanocytes, and endothelial cells, were shared between the two tissues, whereas TM fibroblasts were specific to TM, and ciliary muscle cells as well as CB_PCE/NPCE cells were specific to CB (Fig. 1c; Supplementary Fig. 1). All clusters contained cells from multiple donors of both sexes with diverse ancestry and age ranges, supporting the robustness of the clustering (Supplementary Fig. 1).

These major cell classes were further resolved into 21 distinct cell types, representing a substantial increase compared to previous reports^3, 4, 17^(Fig. 1e; Fig. 3c). For example, both TM and CB fibroblasts were further refined by re-clustering, revealing multiple distinct cell types (Fig. 1f; Fig. 3a, c). Detailed characterization of these fibroblast populations is provided in subsequent sections. Moreover, epithelial populations in CB were further divided into pigmented and non-pigmented ciliary epithelium (PCE and NPCE) each subdivided into two distinct clusters (PCE_1/2 and NPCE_1/2), which exhibited distinct transcriptomic profiles and clustering characteristics.

A Similar process has been applied to scRNA-seq dataset, resulting in the same nine cell classes, which can be further divided into 19 cell types (Supplementary Fig. 2). Due to the cell number limitation, CB fibroblast from scRNA-seq dataset cannot well-divided into several cell types like snRNA dataset. All clusters comprised cells from multiple donors of both sexes and spanning diverse ancestry groups and age ranges, supporting the robustness of the clustering (Supplementary Fig. 2). For integrated analysis, single-cell variational inference (scVI)^18^was used to construct the atlas based on a benchmark comparison of multiple tools (Supplementary Fig. 3). To assess the consistency between snRNA and scRNA datasets, cell-type proportions were calculated and compared between TM and CB tissues (Fig. 1g). Stromal populations, including ciliary muscle cells (20.39% vs 3.62%), CB_PCE (23.33% vs 2.86%), and CB_NPCE (14.55% vs 1.83%), were substantially enriched in snRNA-seq datasets. In contrast, scRNA-seq datasets showed higher proportions of fibroblasts (32.82% vs 18.88%), Schwann cells (26.03% vs 8.26%), and immune cells (15.39% vs 5.32%). Endothelial cells and pericytes were also relatively enriched in scRNA-seq compared to snRNA-seq. Melanocytes exhibited comparable proportions between the two technologies (6.49% vs 6.33%; Fig. 1g). Cell clusters derived from the two technologies were readily aligned, as they shared consistent transcriptomic signatures across major cell classes. Independent scRNA-seq dataset embeddings further recapitulated the principal transcriptional identities defined in the snRNA dataset analysis, supporting robustness across platforms (Fig. 1h, Supplementary Fig. 3). Further sub-clustering of the major cell classes resolved about 19 distinct cell types, including diverse populations of ciliary epithelial cells, fibroblasts, endothelial cells, Schwann cells, and immune cells (Fig. 1i, Supplementary Fig. 2). Overall, these results demonstrate that, despite differences in cell-type composition, snRNA-seq and scRNA-seq provide complementary and concordant representations of TM and CB cellular architecture.

### Xenium transcriptomics provides spatial evidence for the HTCCA

To provide spatial context for the HTCCA, we performed Xenium in situ spatial transcriptomics on formalin-fixed paraffin-embedded (FFPE) sections of human anterior segment tissues from one donor using a custom designed probe set (Supplementary Table 2). Transcripts were assigned to individual cells based on nuclear and transcript density–guided segmentation, generating a spatially resolved gene expression matrix. Following data acquisition, the full anterior segment dataset was processed in Xenium Explorer v3, from which TM and CB regions were identified for subsequent analyses (Fig. 2a). Within the endothelial cell class, we annotated two distinct endothelial populations, including Schlemm endothelium and vascular endothelium. Notably, Schlemm endothelial cells were exclusively identified within TM samples, whereas vascular endothelial cells were detected across both TM and CB regions (Fig. 1e, f; Supplementary Fig. 1f). Integration with the snRNA reference atlas enabled robust annotation of spatially resolved cell populations. Co-embedding of snRNA-seq and Xenium datasets was performed using Harmony based on shared gene features, allowing alignment of the two modalities in a shared low-dimensional space. UMAP visualization demonstrated clear separation of cell types in both TM and CB, including individual cell types of TM/CB fibroblasts, endothelial cells, pericytes, Schwann cells, melanocytes, immune cells, and CB epithelial populations (CB_PCE and CB_NPCE) (Fig. 2b, f). Cell-type composition across individual slides was largely consistent, with reproducible proportions observed for major cell classes in both CB and TM samples (Fig. 2e, h). Furthermore, heatmaps of marker gene expression in Xenium data further confirmed the identities of annotated cell populations in both CB and TM, demonstrating strong concordance with the RNA atlas (Fig. 2d, i). To further examine spatial changes in cellular composition across the TM, we quantified cell-type proportions along the outer-to-inner axis. This analysis revealed a gradual shift in cellular composition across the TM tissue, with distinct cell populations showing region-dependent enrichment from the outer to inner TM. TM_1 cells were more frequent in the posterior juxtacanalicular region near the SC lumen, whereas TM_3 cells were overrepresented in the anterior uveoscleral region close to the anterior chamber. In contrast, TM_2 cells were relatively evenly distributed in the intermediate region of the TM (Fig. 2j). These results are consistent with previous studies demonstrating three functionally specialized TM cell types with distinct spatial localization patterns in mouse (human cell types were named according to molecular alignment with mouse cell types, while cell type locations were similar across species)^19^.

**Fig. 2.**
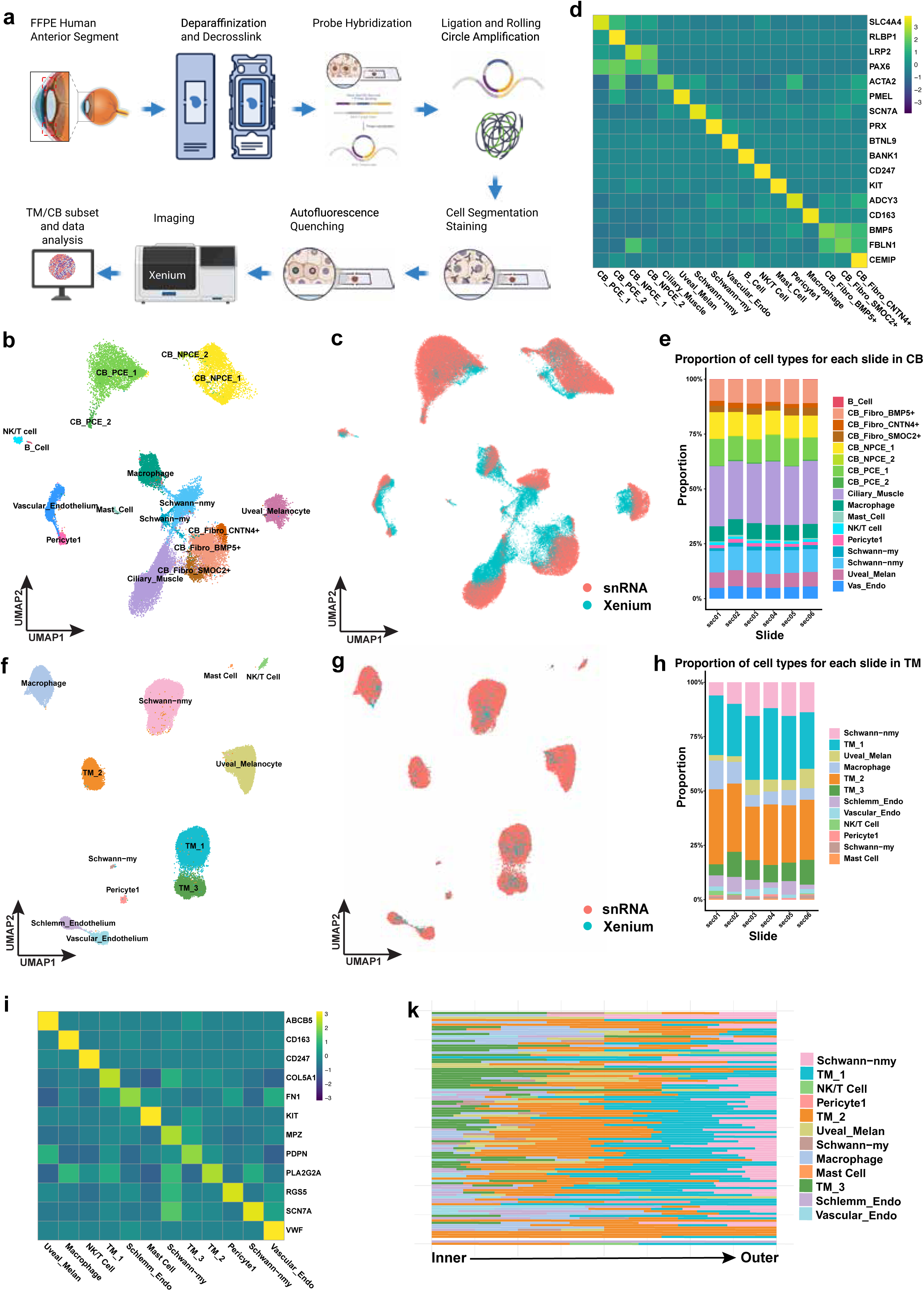
Xenium spatial evidence of the HTCCA. **a**, Schematic overview of the Xenium spatial transcriptomics workflow applied to FFPE sections of the human anterior segment. **b,** UMAP visualization of Xenium and snRNA-seq integrated cells from the CB, with cell types annotated based on the snRNA-seq reference atlas. **c,** UMAP of CB Xenium and snRNA-seq datasets co-embedded using Harmony, colored by dataset origin. **d**, Heatmap of marker genes expression in CB Xenium dataset. **e,** Cell-type composition across individual CB Xenium sections, showing consistent proportions of major cell classes across slides. **f,** UMAP visualization of Xenium and snRNA-seq integrated cells from the TM, with cell types annotated based on the snRNA-seq reference atlas**. g,** UMAP of TM Xenium and snRNA-seq datasets co-embedded using Harmony, colored by dataset origin. **h,** Cell-type composition across individual TM Xenium sections, showing consistent proportions of major cell classes across slides. **i,** Heatmap of marker genes expression in TM Xenium dataset. **j,** Cell-type composition along the outer-to-inner axis of the TM. Each horizontal bin represents a spatial interval across the TM, and stacked bars show the relative proportions of annotated cell types within each bin. Uveal_Melan, Uveal melanocytes; Vas_Endo, Vascular Endothelium.

**Fig. 3.**
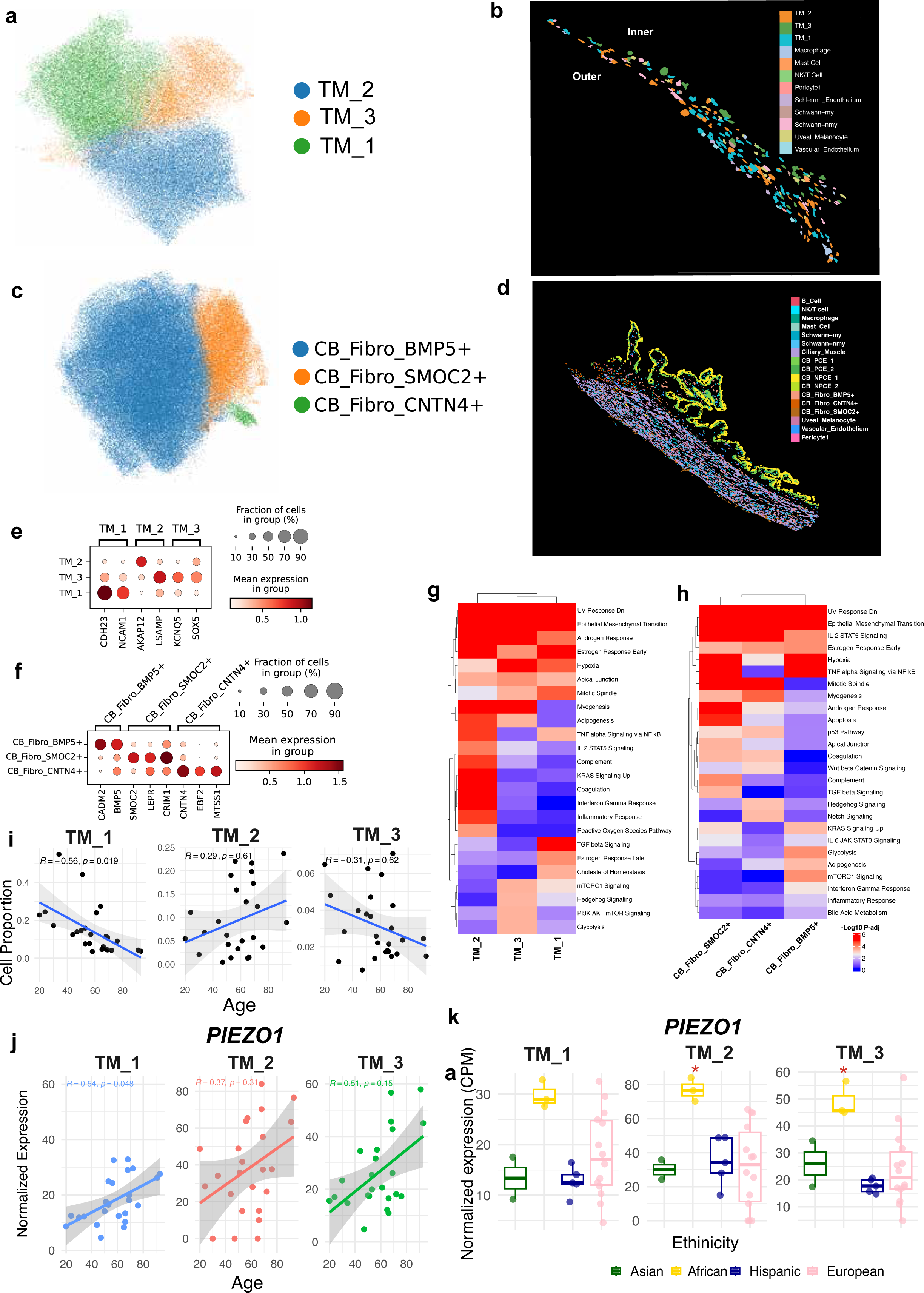
Identification and characterization of TM and CB fibroblast. **a,** UMAP visualization of TM fibroblast cell types. **b,** Spatial localization of TM fibroblast cell types in Xenium data. **c,** UMAP visualization of CB fibroblast cell types. **d,** Spatial localization of TM fibroblast cell types in Xenium data. **e,** Dot plot showing representative marker genes for TM fibroblast cell types. Dot size indicates the fraction of expressing cells and color intensity represents average expression level. **f,** Dot plot showing representative marker genes for CB fibroblast cell types. **g,** Hallmark pathway enrichment analysis of TM fibroblast cell types. **h,** Hallmark pathway enrichment analysis of CB fibroblast cell types. **I,** Correlation between donor age and relative proportions of TM fibroblast cell types across donors. **j,** Correlation between donor age and normalized expression levels of representative cell type associated genes in TM fibroblast populations. **k,** Comparison of TM fibroblast cell type proportions across different ancestry groups. Boxplots represent normalized expression or cell proportion distributions across donors from different ancestries.

In the CB, consistent with known CB anatomy, PCE and NPCE cells form well-defined, closely opposed layers arranged in an apex-to-apex orientation^4^(Fig. 3d). These epithelial domains were spatially confined and clearly separated from the underlying stromal compartment. The stromal region was enriched for fibroblasts, vascular endothelial cells, pericytes, and immune populations, consistent with its role as a vascularized connective tissue core (Fig. 3d). Ciliary muscle cells were localized adjacent to the stromal compartment, forming a distinct muscular region (Fig. 3d). In addition, melanocytes and Schwann cells exhibited region-specific distributions within the CB, further highlighting the spatial heterogeneity of this tissue (Fig. 3d). Together, these spatial patterns recapitulate the known structural organization of the TM/CB and support the cell type identities defined by the RNA atlas, demonstrating that Xenium spatial transcriptomics enables single-cell resolution mapping of cellular architecture in human TM and CB tissues.

### Fibroblasts in TM and CB

Fibroblasts represent a major stromal cell population in both the trabecular TM and CB and play essential roles in maintaining tissue structure and function. TM fibroblasts are present across all three anatomical regions of the TM, with those in the juxtacanalicular region adjacent to Schlemm’s canal embedded within a dynamic extracellular matrix and playing a key role in regulating aqueous humor outflow.^9, 19^ Following re-clustering of TM fibroblasts, we identified three distinct sub-clusters. Based on previously reported marker genes^4, 17^, these clusters were previously annotated as JCT cells, Beam A, and Beam B, and were hereafter referred to as TM_1, TM_2, and TM_3, respectively (Fig. 3a, e). In the snRNA-seq atlas, each cell type exhibited distinct marker gene expression profiles. TM_1 cells exhibited high expression of *ANGPTL7*, TM_2 cells were characterized by enrichment of *BMP5*, whereas TM_3 cells preferentially expressed *TMEFF2* (Fig. 1f). Notably, our Xenium spatial transcriptomics revealed clear spatial biases in the distribution of these cell types across the TM. TM_1 cells were preferentially localized to the outer regions of the TM, whereas TM_3 cells were enriched in the inner regions adjacent to SC. TM_2 cells were primarily distributed in intermediate regions of the TM (Fig. 2j; 3b; Supplementary Fig. 7). These spatial patterns are consistent with previous observations in mouse TM^19^, further supporting the robustness and biological relevance of our cell type annotation. Interestingly, TM fibroblast cell types exhibited distinct age- and ancestry-associated patterns (Fig. 3i-k). TM_1 cell proportion significantly decreased with age, whereas TM_1 associated gene *PIEZO1* expression increased with age (Fig. 3i, j); in addition, TM_2 and TM_3 showed higher normalized expression levels of gene *PIEZO1* in African donors compared with other ethnic groups (Fig. 3k).

For fibroblast cells in the CB, three distinct cell types are identified (Fig. 3c). As CB fibroblast heterogeneity has not been well characterized in previous studies, we performed differentially expressed gene (DEG) analysis to identify cell type-specific marker genes. Each cell type exhibited distinct transcriptional signatures, with enrichment of *BMP5*, *SMOC2*, and *CNTN4* genes, respectively, and were therefore designated as CB_Fibro_BMP5+, CB_Fibro_SMOC2+, and CB_Fibro_CNTN4+ (Fig. 3c, f). Notably, Xenium spatial analysis showed that, unlike TM fibroblasts, these CB fibroblast cell types were broadly distributed across the stromal region, without clear spatial segregation or positional preference.

Hallmark gene set enrichment analysis revealed distinct functional programs among TM and CB fibroblast cell types (Fig. 3g, h). TM_1 cells were enriched for metabolic and TGF-β–related pathways, TM_2 preferentially enriched with EMT, hypoxia, and apical junction pathways, whereas TM_3 showed enrichment of inflammatory and stress-response pathways, including TNFα/NF-κB and interferon gamma signaling (Fig. 3g). Similarly, CB fibroblast cell types displayed distinct transcriptional signatures, with CB_Fibro_BMP5+ enriched for IL6/JAK/STAT3 signaling and glycolysis, CB_Fibro_SMOC2+ enriched for EMT, TGF-β, and Hedgehog signaling, and CB_Fibro_CNTN4+ enriched for inflammatory-related pathways (Fig. 3h).

### HTCCA: chromatin accessibility landscape

To understand how gene regulation shapes different TM and CB cell classes and types, we profiled chromatin accessibility landscapes of TM and CB cells using snATAC-seq. After quality control, we obtained 725,665 high-quality nuclei. We annotated 9 major classes in the snATAC-seq dataset by co-embedding it with snRNA-seq data using scGLUE^20^. These snATAC-seq cell classes aligned with the ones identified in the snRNA-seq data (Fig. 4a). The overall cell composition was highly consistent between snRNA-seq and snATAC-seq datasets, with CB_PCE as the most abundant cell class, followed by ciliary muscle, fibroblasts, CP_NPCE, immune cells, Schwann cells, melanocytes, endothelial cells and pericytes (Fig. 4a, b). Canonical marker genes showed cell class-specific chromatin accessibility and inferred gene activity patterns that mirrored their expression profiles in the snRNA-seq data (Fig. 4c).

**Fig. 4.**
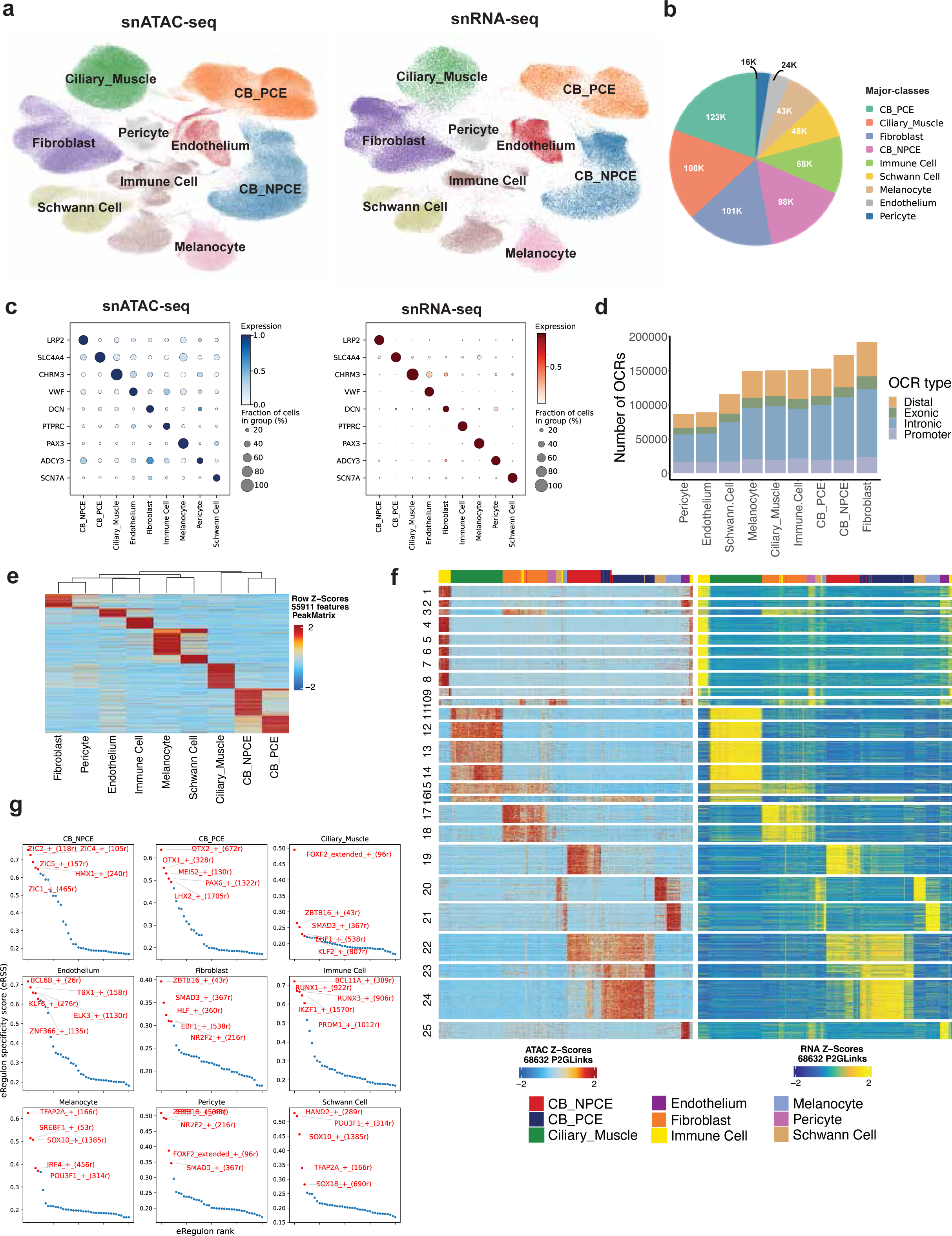
A high resolution snATAC-seq cell atlas of the human TM and CB. **a,** UMAP of co-embedded cells from snATAC-seq (left) and snRNA-seq (right) showing cells are clustered into major cell classes. **b,** Pie chart showing the cell proportion distribution of major cell classes. **c,** Dot plot showing marker gene expression measured by snRNA-seq and marker gene activity score derived from snATAC-seq in the corresponding cell class. **d,** Bar plot showing the number of open chromatin regions (OCRs) identified in each major cell class. **e,** Heatmap showing the chromatin accessibility of differential accessible regions (DARs) identified in major retinal cell classes. Rows represented regions and columns corresponded to cell classes. **f,** Heatmap showing chromatin accessibility (left) and gene expression (right) of 68,632 significantly linked CRE-gene pairs identified by the correlation between gene expression and OCR accessibility. Rows represent cis-regulatory element (CRE)-gene pairs, which are clustered into 25 groups using k-means clustering. Columns represent cell groups that were grouped using the K Nearest Neighbor (KNN) method. **g,** Scatter plot showing eRegulon specificity score of each transcription factor (TF) regulon across major classes. The top five TF are highlighted in red.

From this dataset, we identified open chromatin regions (OCRs) in each cell class, ranging from 86,528 OCRs in pericytes to 191,740 OCRs in fibroblasts (Fig. 4d). As expected, the majority of OCRs were located in intronic regions, followed by distal regions, promoters, and exonic regions (Fig. 4d). While many OCRs were shared across multiple cell classes, 0.69% to 7.19% per class displayed cell class–specific accessibility. These regions likely contribute to defining cell-type–specific gene regulation and are referred to as differentially accessible regions (DARs) (Fig. 4e). To link OCRs to their potential target genes, we examined correlation between gene expression or promoter accessibility and nearby OCR accessibility within -/+ 250 kb. This analysis identified 68,633 OCR-gene pairs (Fig. 4f), representing putative cis-regulatory elements and their likely target genes.

We next identified TFs that define major TM and CB cell classes by integrating snRNA-seq and snATAC-seq data using SCENIC+^21^ (Fig. 4g, Supplementary Fig. 8a, Supplementary table 6, 7). Many of the TFs identified are well-established regulators of TM and CB cell identity, including EBF1 and SMAD3 in fibroblasts, ZIC family members and HMX1 in CB_NPCE, OTX1/OTX2 and PAX6 in CB_PCE, FOXP2, SMAD3 and EBF1 in ciliary muscle cells, BCL6B in endothelial cells and ERF1 in Pericytes (Fig. 4g). Most interestingly, in fibroblasts, regulatory modules involving SMAD3, EBF1, and NR2F2 were prominently enriched, suggesting active roles of TGF-β–related and transcriptional regulatory networks. These findings highlight the presence of cell type–specific gene regulatory architectures underlying functional specialization in anterior segment tissues. In addition, we uncovered candidate TFs not previously linked to specific TM and CB cell classes, providing new insights into regulatory networks governing these tissues. Correlation analysis of regulon activity further demonstrated structured relationships among transcriptional programs, with cell type–specific clusters of regulons reflecting coordinated regulatory networks underlying cellular identity and function, as evidenced by shared cis-regulatory regions and targeted genes (Supplementary Fig. 8b, c).

To further resolve cell types within each class, we co-embedded snATAC-seq and snRNA-seq datasets using scGLUE and applied a logistic regression model trained on snRNA-seq data to predict cell types in snATAC-seq data (Supplementary 9c). This approach identified 19 cell types, matching the 19 types defined by snRNA-seq (Supplementary Fig. 9a, b).

### Age- and ancestry-related changes in TM and CB cell populations

The broad age range of our dataset enables us to investigate age-related molecular and cellular changes in the TM and CB cell populations. Remarkably, we observed significant age-associated changes in cell proportions. In the CB, after controlling potential confounders, such as ancestry and sex, endothelial cells (*p*=2.4e-4), fibroblasts (*p*=0.029), immune cells (*p*=0.0088) were significantly reduced with age, whereas Schwann cells (p=0.036) were significantly increased (Fig. 5a). In the TM, endothelial cells (*p*=0.057) and fibroblasts (*p*=0.082) exhibited marginal decrease with age, while Schwann cells (*p*=0.023) similarly increased with age. In contrast, immune cells did not show a significant change with age (Fig. 5b).

**Fig. 5.**
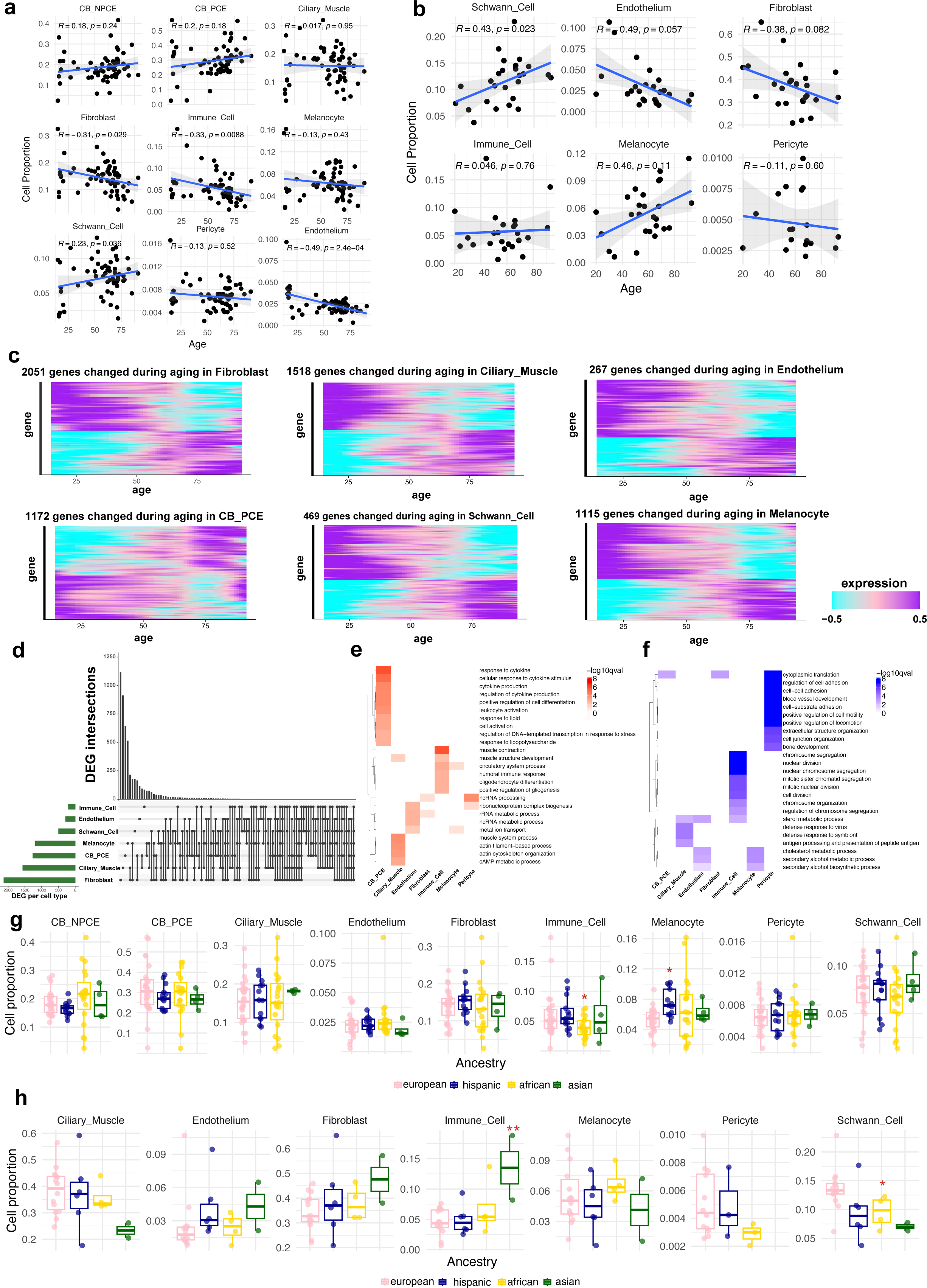
Changes of TM and CB cell populations associated with age and ancestry. **a,** Scatter plots showing age-associated changes in cell proportions in the CB. The X-axis indicates age (years). The Y-axis indicates cell proportions. Each dot represents one sample. Pearson correlation coefficients R and P-values from multivariable linear regression (adjusted for ancestry and sex) are showed in the plots. **b,** Scatter plots showing age-associated changes in cell proportions in the TM. The X-axis indicates age (years). The Y-axis indicates cell proportions. Each dot represents one sample. Pearson correlation coefficients R and *p*-values from multivariable linear regression (adjusted for ancestry and sex) are showed in the plots. **c,** Heatmap showing gene expression of DEGs during aging in major cell classes identified with LMM. **d,** Upset plot showing the numbers of shared and distinct DEGs across major cell classes. **e,** Heatmap showing enriched biological processes among up-regulated DEGs across major cell classes. **f,** Heatmap showing enriched biological processes among down-regulated DEGs across major cell classes. **g,** Box plot showing the distribution of cell proportions across ancestry groups for each major cell class in the CB. Significant P-values are derived from multivariable linear regression (adjusted for age and sex) *: *p* < 0.05, **: *p* < 0.01, and *** *p* < 0.001. **h,** Box plot showing the distribution of cell proportions across ancestry groups for each major cell class in the TM. Significant p-values are derived from multivariable linear regression (adjusted for age and sex).

In addition to changes in cell proportions, we observed pronounced age-associated changes in gene expression. The number of DEGs is up to 2051 across major cell classes, with the highest numbers detected in fibroblasts, followed by ciliary muscle cells, CB_PCE, melanocytes, Schwann cells, endothelial cells, and immune cells (Fig. 5c, supplementary Fig. 9d). Most DEGs were cell class–specific, with a moderate fraction shared across cell classes (Fig. 5d), and were enriched in cell type–specific biological processes and pathways (Fig. 5e, f).

For example, DEGs upregulated with age in CB_PCE were enriched in pathways related to cytokine response, lipid response, and stress responses. In contrast, DEGs upregulated in ciliary muscle cells were enriched in actin filament–based processes and cAMP metabolic pathways (Fig. 5e). Among downregulated genes, pericytes showed enrichment for processes such as cytoplasmic translation, regulation of cell adhesion, blood vessel development, and cell junction organization. Whereas age-downregulated DEGs in immune cells were enriched in cell cycle–related processes, including chromosome segregation and mitotic nuclear division, as well as sterol metabolic processes (Fig. 5f).

Interestingly, cell proportions also exhibited ancestry-related differences. In the CB, immune cells are less abundant in African ancestry compared with European (*p*<0.05, after controlling age and sex). In contrast, melanocytes were more abundant in Hispanic individuals (*p*<0.05, after controlling age and sex) (Fig 5g). In the TM, immune cells are more abundant in Asian ancestry (*p* < 0.01, after controlling age and sex), whereas Schwann cells were less abundant in individuals of African ancestry compared with European (*p*<0.05, after controlling age and sex) (Fig. 5h).

### GWAS cell type enrichment and fine-mapping

We integrated the multimodal HTCCA atlas with GWAS data to systematically prioritize putative causal variants, genes, and cell types relevant to eye diseases and traits. GWAS signal enrichment across cell classes was assessed using complementary approaches, including gene expression-based MAGMA analysis and chromatin accessibility-based LDSC analysis (Fig. 6a, b).

**Fig. 6.**
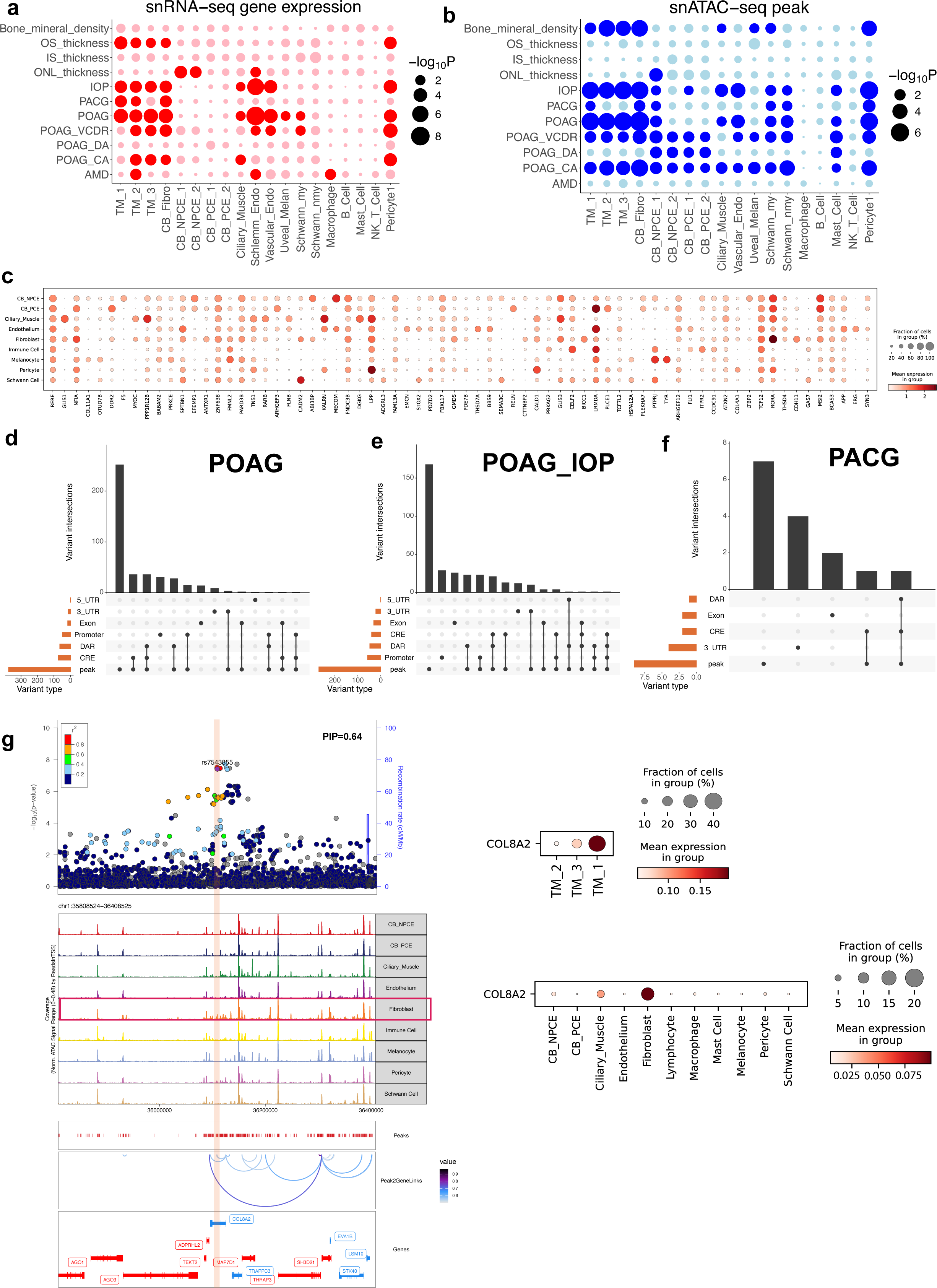
Cell type enrichment and fine-mapping of GWAS loci. **a,** Enrichment of GWAS loci across CB and TM cell types based on snRNA-seq gene expression with MAGMA. Celltyping. Rows indicate GWAS traits, and columns indicate cell types. The dot size indicates −log10P. Significant results (FDR < 0.1) are highlighted in red, while non-significant results (FDR ≥ 0.1) are shown in pink. **b,** Enrichment of GWAS loci across CB and TM cell types based on snATAC-seq chromatin accessibility with LDSC. Rows indicate GWAS traits, and columns indicate cell types. The dot size indicates −log10P. Significant results (FDR < 0.1) are highlighted in blue, while non-significant results (FDR ≥ 0.1) are shown in light blue. **c,** Dotplot showing expression of selected GWAS candidate genes across major cell classes. **d,** Upset plot showing the distribution of genomic annotations among fine-mapped POAG GWAS variants, including peak (OCRs), CREs, DARs, promoter, exon, 5’ and 3’ UTR regions. **e,** Upset plot showing the distribution of genomic annotations among fine-mapped UKBB_IOP GWAS variants. **f,** Upset plot showing the distribution of genomic annotations among fine-mapped PACG GWAS variants. **g,** Visualization of a fine-mapped loci (rs7543855) in *COL8A2* region. Dot plots showing the normalized expression level of *COL8A2* across major cell classes and TM fibroblast cell types. LocusZoom plots showing the GWAS signals. Genome tracks showing the OCRs across cell classes from snATAC–seq data.

Glaucoma-related traits showed cell type-specific enrichment in fibroblasts, ciliary epithelial cells, ciliary muscle cells, pericytes, and endothelial cells (Fig. 6a, b). Specifically, loci associated with primary open-angle glaucoma (POAG) were enriched in TM_1, TM_2, TM_3, CB fibroblasts, pericytes, ciliary muscle cells, Uveal melanocytes, myelinating Schwann cells, Schlemm’s canal endothelial cells, and vascular endothelial cells (Fig. 6a, b; Supplementary Fig. 5b). Traits related to optic disc morphology also showed enrichment in TM and CB cell types: vertical cup-to-disc ratio were enriched in TM_2, TM_3, ciliary fibroblasts, myelinating Schwann cells, SC endothelial cells, vascular endothelial cells, and pericytes. Cup area was enriched in TM_2, TM_3, ciliary fibroblasts, ciliary muscle cells, and pericytes. Optic disc area was enriched in CB non-pigmented and pigmented ciliary epithelial cells (Fig. 6a, b). These data demonstrate the potential correlation between anterior segment tissue and retinal endophenotypes contributing to glaucoma development. In contrast, loci associated with primary angle-closure glaucoma (PACG) were enriched in CB fibroblasts, TM_1, and TM_2 (Fig. 6a, b). To investigate the expression of GWAS candidate genes, we mapped their expression in TM and CB cell classes and types (Fig. 6c). More than 90 glaucoma genes showed high expression across TM and CB cell types (Fig. 6c).

We further fine-mapped GWAS loci associated with POAG, IOP and PACG to prioritize candidate variants^22–24^, genes and cell types (Fig. 6d-f). Based on GWAS summary statistics, 783 variants within 95% credible sets were identified with the ‘Sum of Single Effects’ statistical model. We further categorized fine-mapped variants into functional genomic regions, including HTCCA-derived OCRs, DARs, and CREs, as well as genome annotation-derived promoters, UTRs, and exons. A subset of fine-mapped variants locate in these functional genome regions, the majority of which resided in OCRs (87.2%, 80.1%, and 60.0%), along with notable fractions in CREs (17.2%, 11.9%, and 13.3%) and DARs (15.1%, 14.2%, and 6.7%), supporting their potential regulatory contributions to disease-associated loci (Supplementary Table 8, 9). These fine-mapped variants provide candidates and regulatory hypotheses underlying GWAS loci. Interestingly, we fine-mapped rs7543855, a GWAS variant associated with IOP to a CRE linked to *COL8A2* (PIP = 0.64; Fig. 6g). This CRE is preferentially accessible in fibroblasts and is linked to the expression of *COL8A2* through peak-to-gene correlation analysis (Fig. 6g). Consistent with this regulatory relationship, *COL8A2* is predominantly expressed in fibroblasts. with the highest expression observed in the TM_1 population. In addition, rs7543855 has a high CADD score (8.6) and phyloP score (3.95), supporting its potential functional importance. Taken together, these findings support a cell type-specific regulatory role for this variant in modulating *COL8A2* expression in TM fibroblasts.

## Discussion

The cells of the TM and CB are key components of the anterior segment that produce or drain aqueous humor thereby generating IOP to facilitate vision^7, 9–11, 25–27^. Dysfunction of these tissues, particularly the TM and adjacent SC are major drivers of glaucoma pathogenesis, yet their cellular and regulatory organization in humans remains incompletely understood^11, 14, 28^.

In this study, we constructed a comprehensi ve multi-omics single-cell atlas of the human TM and CB, integrating transcriptomic and chromatin accessibility profiles with spatial transcriptomics. By combining snRNA-seq, scRNA-seq, and snATAC-seq data from over one million cells and nuclei across 112 donors, together with Xenium-based spatial profiling, our HTCCA provides a high-resolution and spatially informed reference of the human anterior segment. This integrated resource captures cellular composition, transcriptional states, regulatory landscapes, and spatial organization, offering a framework for investigating aqueous humor dynamics and anterior segment biology. Compared with previous TM and CB studies, our atlas substantially increases both cellular resolution and sample diversity. We identified 9 major cell classes and further resolved them into 19–21 distinct cell types/cell types, including previously under-characterized fibroblast and epithelial subpopulations. Notably, integration of scRNA-seq and snRNA-seq datasets revealed strong concordance in cell-type identities despite differences in cell-type proportions, highlighting the complementary nature of these technologies.

A key advance of our study is the spatial characterization of cell types and cell types using Xenium transcriptomics. By integrating spatial data with the snRNA-seq reference, we mapped cellular organization at single-cell resolution and confirmed known anatomical features of the TM and CB tissues. In the TM, we observed a clear spatial gradient along the outer-to-inner axis, with TM_1, TM_2, and TM_3 fibroblast cell types occupying distinct yet overlapping zones. These patterns recapitulate prior observations in mouse models and provide direct spatial evidence for cross species conservation in the TM.^7, 19^ In the CB, epithelial populations (PCE and NPCE) formed well defined bilayer structures, while stromal and vascular populations displayed region-specific distributions, reflecting the complex architecture of this tissue.

Fibroblasts emerged as a major source of cellular heterogeneity and functional specialization in both the TM and CB.^3, 4, 7, 15, 17, 28^ In the TM, three fibroblast cell types corresponding to TM_1, TM_2, and TM_3 exhibited distinct transcriptional programs and spatial localization, suggesting specialized roles in extracellular matrix remodeling, mechanotransduction, and immune signaling^27, 29^. In the CB, we identified three fibroblast cell types with distinct molecular signatures but relatively uniform spatial distribution, indicating functional diversification without strong positional segregation. To our knowledge, this represents the first systematic characterization of fibroblast heterogeneity in the human ciliary body at single-cell resolution.

Hallmark pathway analysis further revealed that these fibroblast populations are characterized by distinct biological programs, ranging from metabolic and TGF-β signaling to inflammatory and stress-response pathways. Notably, TGF-β signaling was enriched in TM_1 and CB_Fibro_SMOC2+ populations, suggesting a shared regulatory program associated with extracellular matrix remodeling and fibrotic responses. This observation is consistent with previous studies demonstrating that elevated TGF-β2 levels in aqueous humor drive extracellular matrix accumulation and increased outflow resistance in the TM, contributing to elevated IOP and glaucoma pathogenesis^29^ Moreover, TGF-β–mediated ECM remodeling represents a key mechanism linking mechanical stress, tissue stiffening, and progressive dysfunction in the anterior segment^25, 27, 30^. These distinct pathway enrichments further raise the possibility that fibroblast cell types reflect different functional states associated with microenvironmental cues and activation programs. The distinct Hallmark pathway enrichments observed across TM fibroblast cell types suggest that these populations may, at least in part, reflect dynamic cellular states associated with fibroblast activation and environmental responses. However, their reproducible clustering across donors and spatially restricted distributions within the TM support the presence of structured heterogeneity beyond purely transient states. Together, these findings suggest that TM fibroblast heterogeneity is more consistent with the presence of stable cell types at this resolution, although contributions from dynamic cellular states cannot be excluded. Further analyses will be required to more precisely delineate these relationships.

At the regulatory level, our snATAC-seq analysis delineates a comprehensive chromatin accessibility landscape of TM and CB tissues. We identified extensive open chromatin regions and cell-type–specific DARs that likely underlie transcriptional diversity across cell populations. Integration with gene expression data enabled the prediction of putative cis-regulatory elements and their target genes, providing insights into the regulatory architecture of anterior segment tissues. In addition, TF analysis uncovered both known regulators and previously unrecognized candidates, suggesting complex regulatory networks governing TM and CB cell identity.

Fibroblast populations exhibit pronounced transcriptional and regulatory diversity, with enrichment of SMAD3-associated regulons and TGF-β signaling, supporting activation of extracellular matrix remodeling and fibrotic programs^7, 24, 27, 28^. These findings are consistent with the established role of TGF-β–mediated ECM accumulation in increasing outflow resistance and IOP, and further suggest that fibroblasts act as key mediators linking mechanical stress, tissue stiffening, and anterior^25, 27, 30^ segment dysfunction. At the same time, CB epithelial populations are characterized by enrichment of developmental TF regulons, including members of the OTX and PAX families, consistent with their established roles in ocular development and maintenance of epithelial identity in the CB^31, 32^. The presence of distinct, cell type specific regulatory programs, together with structured spatial organization and reproducible clustering across donors, indicates that cellular heterogeneity in the TM and CB reflects both stable cell identities and dynamic functional states shaped by local microenvironmental cues. Collectively, these findings support a model in which coordinated transcriptional and regulatory networks govern anterior segment homeostasis, while dysregulation of these programs, particularly within fibroblast populations, may contribute to extracellular matrix remodeling, increased outflow resistance, and ultimately the pathogenesis of glaucoma^25, 27^.

Our DEG analysis reveals substantial age-associated remodeling of both cellular composition and transcriptional programs in TM and CB tissues, highlighting aging as a key driver of anterior segment heterogeneity^26, 33^. Notably, TM fibroblast cell types exhibited distinct age- and ancestry-associated patterns. TM_1 cell proportions declined significantly with age, whereas expression of the mechanosensitive ion channel *PIEZO1* increased with age within this population. This divergence between decreasing cell abundance and increasing gene expression suggests functional reprogramming of TM_1 fibroblasts during aging. Given that *PIEZO1* functions as a key mechanosensitive ion channel that mediates cellular responses to mechanical stimuli such as stretch and shear stress^34, 35^, its upregulation is consistent with enhanced mechanosensitive signaling by remaining TM_1 cells.

Given that *PIEZO1* functions as a key mechanosensitive ion channel that mediates cellular responses to mechanical stimuli such as stretch and shear stress, and regulates cytoskeletal organization and aqueous humor outflow in trabecular meshwork cells^34, 36, 37^, its upregulation is consistent with enhanced mechanosensitive signaling and may reflect adaptive responses to cell/tissue stiffening to maintain trabecular meshwork function and IOP homeostasis. In addition, TM_2 and TM_3 fibroblast cell types exhibited higher *PIEZO1* expression in African ancestry donors, and previous studies have investigated a gain of function *PIEZO1* variant that is common in individuals of African descent in relation to glaucoma-related phenotypes. Although the associations with IOP or disease progression is not statistically significant, it is likely due to insufficient power and the reported trends suggest a potential role in IOP regulation and disease related processes^38^.

A particularly striking observation is the age-associated expansion of Schwann cells in CB. This may reflect compensatory responses to neural stress or degeneration and suggests a role for glial activation in anterior segment remodeling. Together with the observed transcriptional decline in immune and vascular associated programs, these findings point to coordinated changes in neural, stromal, and vascular regulation with aging. Collectively, our results support a model in which aging induces functional reprogramming across multiple cell types, particularly fibroblasts and Schwann cells, leading to coordinated alterations in extracellular matrix regulation, mechanosensitive signaling, and neural homeostasis. These processes may converge to disrupt aqueous humor dynamics and increase susceptibility to glaucoma. It is worth noting that Schwann cells were detected in our TM samples and exhibited age-associated expansion, which may reflect age-associated changes in TM anatomy and associated dissection-related contamination.

In our GWAS fine-mapping analysis, we identified multiple glaucoma-associated variants residing within cell type–specific regulatory elements. Notably, we fine-mapped rs7543855, an IOP-associated GWAS variant^23^ to a fibroblast-specific CRE linked to *COL8A2*. Integration of chromatin accessibility and peak-to-gene linkage analyses demonstrated that this regulatory element is preferentially accessible in fibroblasts and is connected to *COL8A2*, suggesting a potential regulatory mechanism underlying this association.

*COL8A2* encodes the α2 chain of type VIII collagen, a key extracellular matrix component implicated in ocular tissue structure and homeostasis^39^. Consistent with this regulatory relationship, *COL8A2* expression was highly enriched in fibroblasts, particularly within the TM_1 fibroblast population. Given the central role of the TM extracellular matrix in regulating aqueous humor outflow resistance and IOP, altered regulation of *COL8A2* may influence conventional outflow pathway function through effects on extracellular matrix composition and biomechanical properties. Consistently, *COL8A2* has been associated with IOP and POAG in previous GWAS studies^23, 24^.

Importantly, rs7543855 was prioritized with a high posterior inclusion probability (PIP = 0.64), supporting its candidacy as a causal regulatory variant at this locus. The convergence of genetic association, chromatin accessibility, peak-to-gene linkage, and cell type–specific gene expression provides multiple lines of evidence nominating *COL8A2* as a likely target gene of this glaucoma-associated locus. Together, these findings establish a potential mechanistic connection between glaucoma-related trait risk and TM fibroblast biology, highlighting extracellular matrix regulation as a key pathway linking genetic variation to anterior segment dysfunction and elevated IOP. More broadly, these results demonstrate the value of integrating GWAS fine-mapping with single-cell transcriptomic and epigenomic data to assign candidate target genes and cellular contexts to non-coding risk variants. Such approaches provide a framework for translating genetic associations into testable biological mechanisms relevant to glaucoma pathogenesis.

In summary, our HTCCA represents a comprehensive reference of the human TM and CB, capturing cellular, spatial, and regulatory heterogeneity at unprecedented resolution. This resource provides a foundation for future studies of aqueous humor regulation, aging, and glaucoma pathogenesis, and offers a roadmap for integrating multi-modal data to dissect complex ocular diseases. However, while we provide a comprehensive characterization of TM and CB, key components of the conventional outflow pathway, particularly SC and distal vessels, are not fully represented. Building on this dataset, future studies incorporating SC and glaucoma donor samples will provide deeper insights into the mechanisms underlying glaucoma pathogenesis.

## Methods

### Ethics

Our study complies with all relevant ethical regulations and was approved by the institutional review boards at the University of California, Irvine; Baylor College of Medicine and the University of Utah.

### Human TM and CB sample preparation

Human donor eyes were recovered from the Baylor College of Medicine Lions Eye Bank and Utah Lions Eye Bank and processed for anterior segment dissection following the established protocol. Detailed donor information is provided in Supplementary Table 1. Donor medical histories were reviewed, and only individuals without evidence of ocular pathology, including glaucoma, were included in the study. Institutional approval for the patient tissue donation consent was obtained from the University of Utah and Baylor College of Medicine, adhering to the tenets of the Declaration of Helsinki. Each tissue was de-identified in accordance with HIPAA Privacy Rules. Under a dissection microscope, the CB was peeled away and collected, and the TM was carefully dissected from the base of the limbal cornea using fine forceps. As TM tissue was dissected by peeling from the limbal region, the outer wall of SC and distal outflow structures are likely underrepresented in this dataset. Therefore, interpretations related to these compartments are inferred from adjacent cell populations and integrative analyses. Fresh tissues were proceeded with single-cell dissociation directly for scRNA sequencing or transferred immediately into ice-cold MACS Tissue Storage Solution for single-cell dissociation within 24 hours, whereas tissues intended for nuclei isolation were snap-frozen and stored in a -80℃ freezer. Eyes with shorter postmortem intervals were prioritized.

### Single-cell dissociation

For single-cell isolation from fresh human TM and CB tissue, tissues were minced into small pieces and dissociated in collagenase A (Sigma, Cat#COLLA-RO) diluted in HBSS at 37°C with shaking. Based on tissue type, ciliary body tissue was incubated in 1 mg/mL collagenase A for approximately 30–60 min, and TM tissue was incubated in 1.5 mg/mL collagenase A for approximately 1 h. Samples were triturated at regular intervals using a wide-bore tip until most tissue fragments had disappeared. After enzymatic dissociation, suspensions were centrifuged at 500 × g for 5 min. To further dissociate ciliary body epithelium, the supernatant was removed and the pellet was resuspended in 0.25% trypsin-EDTA, incubated at 37°C with intermittent trituration for ∼30 min until a single-cell suspension was obtained. For TM samples, the workflow proceeded directly to DNase I treatment after collagenase digestion. DNase I was then added to a final concentration of 80 KU/mL and incubated for 10 min. Cell suspensions were diluted in DMEM containing 5% FBS, filtered through a pre-wet 40 μm strainer, centrifuged at 500 × g for 8 min, and resuspended in an appropriate buffer for cell counting, viability assessment, and immediate loading onto the 10x Genomics Chromium platform. Because CB tissue is highly pigmented, density-gradient cleanup can be used if needed to reduce melanin carryover, although this was balanced against potential loss of pigmented cell populations.

### Single-nucleus isolation and library preparation

For snRNA-seq, tissues were processed for nuclei isolation using the GentleMACS Octo Dissociator with C-tubes in ice-cold Nuclei Extraction Buffer. The resulting suspension was filtered through 70-µm MACS SmartStrainers and centrifuged at 300 × g for 5 min at 4 °C. For TM samples, the nuclei pellet was resuspended in wash buffer (10 mM Tris–HCl, 10 mM NaCl, 3 mM MgCl₂, 1% BSA), gently triturated, passed through 30-µm filters, and centrifuged again at 500 × g for 5 min at 4 °C. The final pellet was resuspended in PBS for downstream processing. For CB samples, nuclei were further purified using the Anti-Nucleus MicroBeads protocol. Briefly, the pellet was resuspended in nuclei separation buffer consisting of Nuclei Extraction Buffer (14%), MACS BSA Stock Solution (0.04%), and RNase inhibitor (0.2 U/µL) in PBS. The suspension was gently triturated, passed through 30-µm filters, and incubated with 50 µL of Anti-Nucleus MicroBeads for 15 min at 2–8 °C. During incubation, LS Columns were prepared according to the manufacturer’s instructions. After incubation, 2 mL of nuclei separation buffer was added to the sample and applied to the magnetic column. The column was washed, and magnetically labeled nuclei were immediately eluted. The collected nuclei were centrifuged at 500 × g for 5 min at 4 °C and resuspended in PBS for downstream processing. Single-nucleus Gene Expression libraries were generated using the Chromium Next GEM Single Cell 3’ v3.1 Reagent Kit (10x Genomics), and single-nucleus ATAC libraries were prepared using the Chromium Next GEM Single Cell ATAC v1.1 protocol.

### Xenium tissue processing of TM and CB

Human ocular anterior segment tissues containing the TM and CB were processed under standardized conditions to preserve tissue morphology and RNA integrity. Detailed donor information is provided in Supplementary Table 2. To facilitate fixative penetration, small incisions were introduced at the pars plana prior to immersion fixation. Tissues were fixed for 48 hours using either 4% paraformaldehyde (PFA) supplemented with 5% sucrose or Modified Davidson’s Fixative, depending on the donor sample. Following fixation, tissues were briefly stored in 1% PFA/5% sucrose or 70% ethanol prior to paraffin embedding. Samples were dehydrated through a graded ethanol series, cleared in HistoClear, and infiltrated with molten paraffin. Paraffin blocks were trimmed to enrich anterior segment regions, and stored at −20°C until sectioning.

### Xenium spatial transcriptomic profiling of TM and CB

Following the 10x Genomics protocol (CG000578_RevE), FFPE tissue blocks were sectioned at 5 μm thickness and mounted onto Xenium slides. Sections were deparaffinized and subjected to RNA decrosslinking according to the manufacturer’s protocol. Spatial transcriptomic profiling was performed using a customized Xenium Human Eye panel targeting 478 genes (Design ID: G33B7Z), with probe hybridization carried out for 16–24 hours.

Signal amplification was achieved through enzymatic ligation and rolling circle amplification, followed by cell segmentation staining and background fluorescence quenching. Imaging and transcript decoding were performed on the Xenium Analyzer, generating cell-by-gene expression matrices and spatial coordinates for each detected transcript. Post-Xenium hematoxylin and eosin (H&E) staining was performed on the same sections to validate tissue morphology and support quality control and segmentation refinement.

### Data collection and preprocessing of snRNA-seq and scRNA-seq data

To achieve the most comprehensive atlas for TM and CB, we applied a similar approach for data collection and processing as used in our human retina cell atlas^5^. Specifically, to collect datasets, we complemented our in-house generated data with public datasets from landmark studies. Raw sequencing reads were downloaded from public data repositories, including the Gene Expression Omnibus (GEO) and Sequence Read Archive (SRA) (Supplementary Table 3). Together with our in-house generated raw sequencing reads, we applied 10x Genomics Cell Ranger v 7.0.1^40^ to generate quantification measurements and used the CellQC^5, 41^ (https://github.com/lijinbio/cellqc; v 0.0.6) pipeline to preprocess the data, including filtering empty droplets using EmptyDrops^43^, correcting ambient RNA contamination using SoupX v 1.6.2^42^, and estimating and filtering doublets using DoubletFinder v 2.0.6^43^. Low-quality cells with <300 features, <500 transcripts, or >10% mitochondrial reads for scRNA-seq (or >5% for snRNA-seq) were filtered out.

### Data integration of scRNA-seq and snRNA-seq and cell clustering

We used scVI v 1.4.2^44^ to integrate scRNA-seq and snRNA-seq datasets. Cell-by-gene matrices from all tissue samples were concatenated using the Scanpy Python package^45^. To capture the variability across the large number of cells, the top 10,000 highly variable genes were first identified using the Scanpy package with the parameters: n_top_genes = 10000, flavor = “seurat_v3”, and batch_key = “sampleID”. Raw counts without normalization for the subsetted highly variable genes were used to fit scVI models with the parameters: batch_key = “sampleID”, n_layers = 2, n_latent = 30, and max_epochs = None. The trained latent representations were used to generate UMAP visualizations. These latent representations were also used for cell clustering using the Leiden algorithm^46^.

To capture the hierarchical nature of cell relationships, we applied a similar two-level clustering approach from the human retina cell atlas^5^ to optimize the resolution parameter. In the first level, various resolution values ranging from 0.1 to 1.0 were tested, and the maximum resolution without over-clustering was selected based on UMAP visualization. For clusters showing compact substructures, a second-level clustering analysis was performed by subsetting the cells and applying another round of Leiden clustering while testing additional resolution values (e.g., ranging from 0.01 to 0.09). The optimal resolution values were selected to achieve well-resolved clusters.

### Integration benchmarking

Benchmarking of integration methods was conducted using an adapted integration workflow from scAtlasTb (https://github.com/HCA-integration/scAtlasTb). The “local” environment mode was selected to install the dependencies, and the required Conda environments were installed manually before running the Snakemake workflow. Three integration methods, scVI^44^, scANVI^47^, and Harmony (Harmonypy)^48^, which are suitable for atlas-scale data integration, were evaluated using default parameters. Each method was configured using 2,000 highly variable genes, sampleID as the batch key, and majorclass/celltype as the label key. Essential evaluation metrics were derived from scIB^18^ and used to evaluate the benchmarked methods. These metrics can be divided into two categories: (1) batch effect removal and (2) biological variance conservation. Metrics in the first category included the average silhouette width (ASW)^48^ batch, graph connectivity, and graph iLISI. Metrics in the second category included ARI, ASW (label), cell cycle conservation, graph cLISI^49^, isolated label ASW, isolated label F1, and NMI. Based on these metrics, individual batch correction and biological conservation scores were calculated for each method. The methods were ranked based on the average of the batch correction and biological conservation scores, referred to as the overall score.

### Cell type annotation and Human-in-the-Loop curation

To annotate cell types, the optimized cell clusters were assigned using known marker genes for major cell classes. Dot plot visualizations were generated to visualize marker gene expression distributions across cell clusters using the Scanpy Python package^45^. Cell clusters that highly and specifically expressed marker genes were annotated to the corresponding major classes. For cross-comparison, existing original cell type labels from public datasets were compared with the identified cell clusters using contingency tables (Supplementary Table 4), and the original labels were used to assist annotation of the major classes. For multiple clusters within an individual major class, suffixes such as “_1” or “_2” were used to distinguish the cell type labels. To achieve robust cell type clustering, annotation, and consensus nomenclature for the atlas, we engaged human-in-the-loop (HITL) community efforts by recruiting voluntary working groups for TM and CB annotation curation using the Cell Annotation Platform (CAP) (https://celltype.info) under the HCA framework. The working group comprised experts with diverse expertise, including physiology, single-cell biology, cross-species analysis, and functional studies. The working group met regularly on a monthly basis to refine clustering, curate annotations, and discuss nomenclature.

### Comparison between scRNA-seq and snRNA-seq

Due to differences in dissociation technologies, we built individual single-cell references for the scRNA-seq and snRNA-seq datasets separately. To compare the two references generated from these technologies, we examined major class similarities using the raw count matrices with the MetaNeighbor R package^50^. The calculated Area Under the Receiver Operating Characteristic Curve (AUROC) represented the similarity across major classes between the two technologies. To examine the two technologies at single-cell resolution, we applied sysVI^51^, which trains a co-embedding between the two references with substantial batches from the two systems corresponding to the two dissociation technologies. We followed the tutorial at https://github.com/Hrovatin/scvi-tutorials/blob/stable/scrna/sysVI.ipynb. The trained latent representation was used to calculate UMAP visualizations of cells from the two technologies.

### Spatial landscape of different cell types using Xenium

The Xenium Onboard Analysis pipeline (version xenium-3.2.0.7, 10x Genomics) was used to generate Xenium bundles for each sample, including four-channel morphology images, decoded transcripts for predefined genes, and negative controls. The four morphology channels comprised DAPI nuclear staining, a cell boundary stain (ATP1A1/E-cadherin/CD45), an interior RNA stain (18S rRNA), and an intracellular protein stain (alphaSMA/Vimentin). Cell segmentation was performed using the multimodal segmentation algorithm implemented in the Xenium Onboard Analysis pipeline. Briefly, nucleus segmentation was first conducted using the DAPI channel, followed by whole-cell segmentation based on boundary and interior staining signals.

Spatial transcriptomic profiling of the TM and CB was conducted using Xenium Ranger (v3, 10x Genomics) with a customized human eye panel comprising 478 genes. Anatomical regions corresponding to TM and CB were delineated based on post-Xenium H&E staining and manually annotated for downstream analyses. The Xenium output was imported into Seurat v5^52^ using the LoadXenium() function for downstream processing^53^. Cells with fewer than 20 detected transcripts were excluded to remove low-quality profiles. The remaining data were normalized and scaled, followed by dimensionality reduction using principal component analysis (PCA). To account for sample-to-sample variability, datasets were integrated using a reciprocal PCA (RPCA)-based approach. The integrated data were embedded in a low-dimensional space using UMAP, and a shared nearest neighbor (SNN) graph was constructed for community detection. Clustering was performed using a modularity optimization algorithm, and cell populations, including major classes and finer cell types within TM and CB, were annotated based on established marker gene expression patterns.

### Processing snATAC-seq and cell type annotation

Raw sequencing reads of snATAC-seq datasets were only collected from in-house generated data (Supplementary Table S5). Raw sequencing reads were processed using the Cell Ranger ATAC pipeline from 10x Genomics (version cellranger-atac-2.0.0). The ArchR package^54^ was used for preprocessing and quality control, including doublet estimation and removal, as well as filtering low-quality cells with transcription start site (TSS) enrichment scores <4 or fewer than 1,000 fragments per cell. To annotate snATAC-seq cells, the snRNA-seq reference was leveraged for label transfer using scGLUE^20^. First, a union reproducible peak set was generated across ATAC cell clusters. Unsupervised integration was then performed to train a co-embedding between snATAC-seq and snRNA-seq cells using cell-by-union peak region count matrices and cell-by-gene count matrices, respectively. The trained co-embedding was subsequently used to train a random forest classifier based on snRNA-seq labels, which was then applied to predict cell type labels for snATAC-seq cells. Following cell type annotation, additional union reproducible peak sets were generated for each cell type for downstream analysis.

### Identification of regulons of different cell types

To identify TFs regulons for different cell types, we applied the SCENIC+ pipeline v1.0a2^21^ using the annotated snRNA-seq and snATAC-seq references. We followed the tutorial (https://scenicplus.readthedocs.io/en/latest/human_cerebellum.html). To efficiently run the analysis workflow, both references were subsampled to a maximum of 2,000 cells per cell type. The precomputed union reproducible peak sets for cell types derived from snATAC-seq were preprocessed using the pycisTopic package. Specifically, cells and union peaks were clustered into regulatory topics using the run_cgs_models_mallet() function, which implements topic modeling using Latent Dirichlet Allocation (LDA)^55^ and runs in Mallet^58^ (Available at: http://mallet.cs.umass.edu). Various topic numbers, ranging from 10 to 150, were tested and evaluated using the evaluate_models() function with “select_model = None”. The optimal topic models were used to extract topic regions using the binarize_topics() function and to identify differentially accessible regions for cell types using the find_diff_features() function. These region sets were subsequently used in the SCENIC+ pipeline to derive eRegulons and their enrichment scores. Downstream gene-target enrichment heatmaps were generated using the heatmap_dotplot() function, and regulon specificity scores were calculated using the plot_rss() function.

### Differential gene expression analysis during aging

To minimize potential biases arising from technological differences between snRNA-seq and scRNA-seq platforms, differential gene expression analyses were performed exclusively on the snRNA-seq dataset. Raw UMI counts were aggregated per gene per sample for each cell class, as most of the samples were profiled from TM and CB regions of the same donors. Samples with lower number of cells within the corresponding cell class were filtered out for downstream analyses. Additionally, samples with gene expression profile (CPM) correlations below 0.75 with more than 65% of other samples were considered as outliers and excluded. The lowly expressed genes were also filtered out. After quality control in both sample and gene levels, genes associated with aging were identified using linear mixed models (LMM) through the edgeR and variancePartition R packages. Gene expression values were normalized using the trimmed mean of M-values (TMM) method. The LMM adjusted for sex, reported ancestry, tissue region, and library size as fixed-effects covariates, and included batch effects as a random intercept.

### Cell type enrichment analysis of GWAS loci

Cell class enrichment of GWAS loci was identified using both chromatin accessibility (snATAC-seq) and gene expression (snRNA-seq) datasets. For gene expression, we assessed linear positive correlations between cell type specificity of gene expression and gene-level genetic association of GWAS studies with MAGMA.Celltype R package. By considering SNPs in - 35kb/+10kb of each gene and 1000G population, the “MungeSumstats” R package was used to format GWAS summary statistics. The “EWCE” R package was employed to format snRNA-seq expression data. We applied MAGMA.Celltype R package to assess linear enrichment. For chromatin accessibility, we use LDSC to partition the heritability of GWAS traits into cell type-specific snATAC-seq peaks with stratified LD score regression. The pipeline annotated GWAS SNPs that overlapped with HapMap3 SNPs based on their location in OCRs in each cell class. 1000G data were employed to calculate the LD scores of these SNPs within 1-cM windows. The LD scores from the baseline model (which included noncell-type-specific annotation, 1000G_Phase3_baselineLD_v2.2_ldscores, downloaded from https://alkesgroup.broadinstitute.org/LDSCORE/) were integrated with the LD scores of these SNPs. To assess whether regions in each cell type were enriched with heritability of the corresponding GWAS trait, the heritability in the annotated genomic regions were estimated and compared with the baseline model. The Benjamini-Hochberg method was applied to the enrichment *P* value from LDSC and MAGMA. Celltyping per trait, accounting for the number of cell types, for multiple-testing correction.

### Fine-mapping of GWAS variants

Initially, statistical fine-mapping of GWAS loci was conducted based on summary statistics from three published glaucoma and IOP GWAS studies^22–24^. For each GWAS study, the SNPs with P < 5×10^−8^ and present in 1000G population were grouped into LD blocks identified by a previous study. The credible set of SNPs and the posterior inclusion probability of each SNP were computed for each LD block, with the “Sum of Single Effects” model via the susieR package (L=10). Subsequently, we integrated genomic annotations, including OCRs, DARs and CREs identified from HTCCA, to annotate and prioritize the fine-mapped variants.

## Data availability

The landing page of the HTCCA data resources is accessible at https://rchenlab.github.io/resources/human-atlas.html. The public datasets used in HTCCA are summarized in Supplementary Table 3. The raw sequencing reads of the newly generated data have been deposited in GEO under accessions GSE326134. Additionally, raw and normalized count matrices, cell-type annotations and embeddings are also publicly available through the CELLxGENE collection (https://cellxgene.cziscience.com/collections/0cbf8ef8-87bd-4b51-8521-ec3f68183e11).

## Code availability

All code used for the HTCCA project can be found in the Github repository (https://github.com/RCHENLAB/HRCA_reproducibility) and (https://github.com/RCHENLAB/HTCCA_reproducibility).

## Author contributions

J.J, J.W. and R.C. conceptualized and designed the study. R.C. supervised the work. X.B. and I.Y collected the samples. Y.Z. performed the Xenium experiments. Y.L. generated the snRNA-seq, scRNA-seq and snATAC–seq data in this study. J.J. and J.W. performed all data analyses. Jin Li provided input to various analysis methods. Jean Li performed the benchmark study analysis. J.J and J.W. wrote the first draft of the paper. All authors edited the paper and contributed to the critical revisions of the paper.

## Acknowledgements

This project was funded by the Chan Zuckerberg Initiative (CZI) 2019-002425 and 2021-239847 to R.C. This work was also supported by NIH grants EY022359 and EY028608 to W.D.S. and NIH grants R01EY025643 to Y.D. National Eye Institute grant EY032507 (to S.W.M.J) the Precision Medicine Initiative at Columbia University, and the New York Fund for Innovation in Research and Scientific Talent (NYFIRST; EMPIRE CU19-2660). The Glaucoma Foundation [NT1] (Columbia University) and an unrestricted departmental award from Research to Prevent Blindness (Columbia University). Additionally, next-generation sequencing (NGS) was performed on instrument supported by the National Institutes of Health (NIH) shared instrument grant S10OD023469 to R.C. The authors acknowledge support to the Gavin Herbert Eye Institute at the University of California, Irvine from an unrestricted grant from Research to Prevent Blindness and from NIH core grant P30 EY034070. We thank Mary Futey, Evan Biederstedt and CAP team for their valuable support and technical assistance. We thank Jennifer Zamanian and the Lattice team at Stanford for their support with data dissemination. This publication is part of the Human Cell Atlas (http://www.humancellatlas.org/publications/). We would also like to acknowledge the feedback and discussions from the members of the HCA Eye Biological Network.

## Competing interests

The authors declare no competing interests.

## Notes

### Competing Interest Statement

The authors have declared no competing interest.

