## Supplementary Figures for "An Integrated muti-omics cell atlas of the human trabecular meshwork and ciliary body"

Supplementary Fig. 1

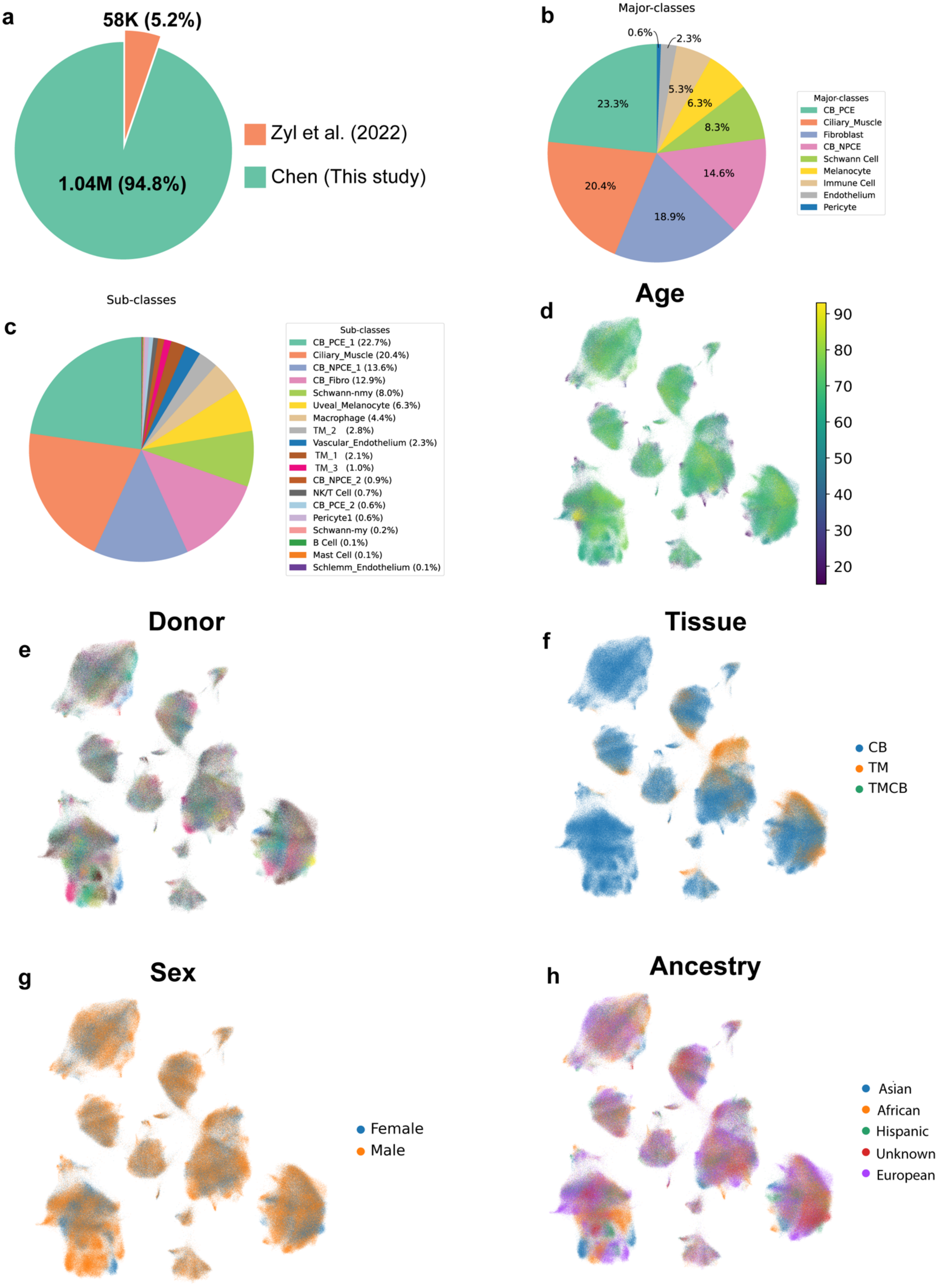

**Supplementary Fig. 1 Overview of dataset composition, cell-type distribution, and donor metadata in snRNA atlas of the human TM and CB**

**a,** Pie charts show the proportion of cells contributed by different study. **b,** Pie charts show the proportion of cells contributed by different study. **b,** Pie charts show the proportion of cells contributed by different major classes. **c,** Dot plots illustrating the distribution of expression levels of marker genes for sub-classes. **d,** UMAP visualization of all cells colored by age. **e,** UMAP visualization of all cells colored by donor. **f,** UMAP visualization of all cells colored by tissue. **g,** UMAP visualization of all cells colored by sex. **h,** UMAP visualization of all cells colored by ancestry.

**Supplementary Fig. 2**

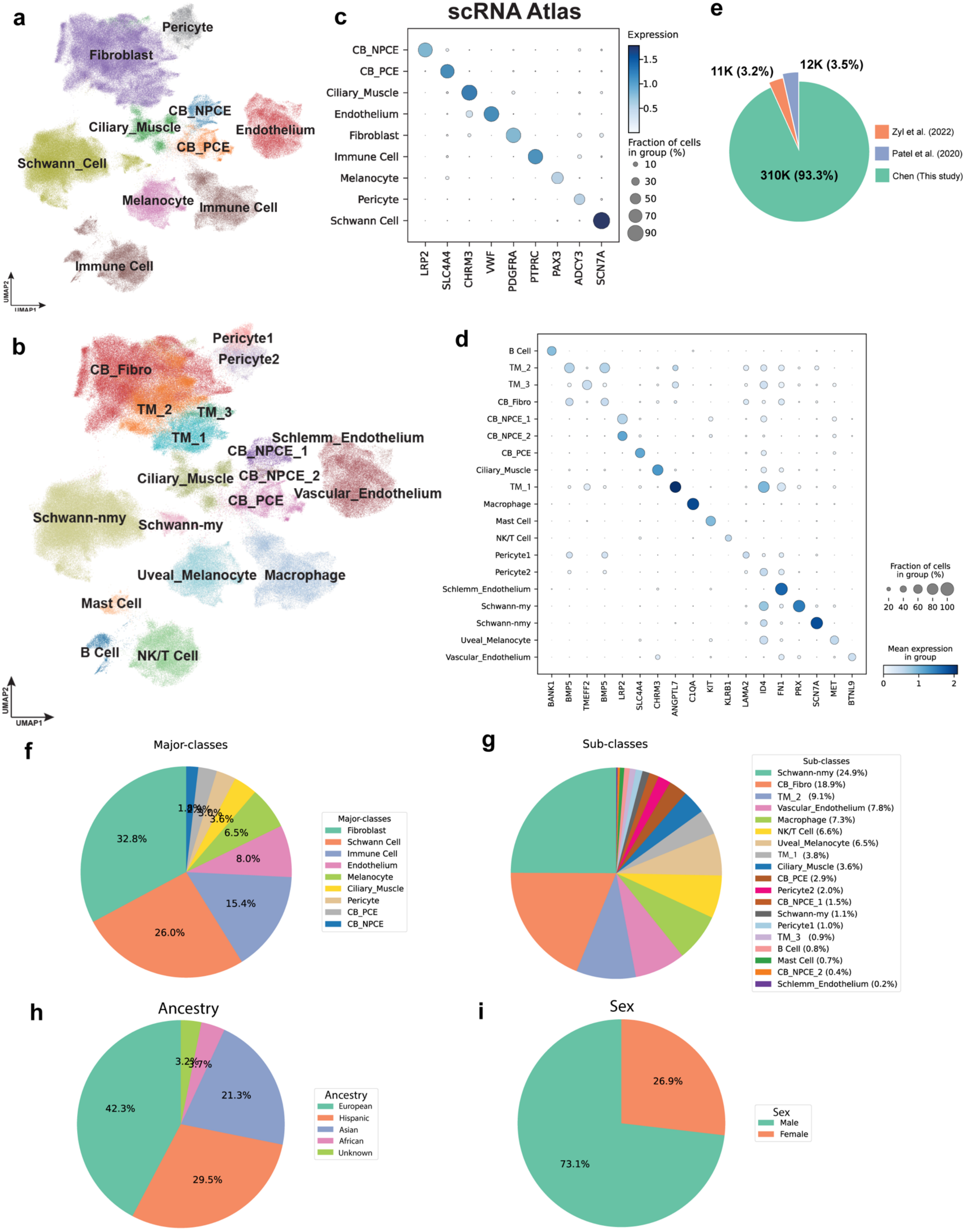

### Supplementary Fig. 2 scRNA atlas of the human TM and CB

**a,** The atlas of scRNA-seq datasets is visualized in a UMAP plot at a major class resolution, with cells colored based on their major classes. **b,** The atlas of scRNA-seq datasets is visualized in a UMAP plot at a major class resolution, with cells colored based on their sub-classes. **c,** Dot plots illustrating the distribution of expression levels of marker genes for major cell classes. **d,** Dot plots illustrating the distribution of expression levels of marker genes for sub-classes. **e,** Pie charts show the proportion of cells contributed by different study. **f,** Pie charts show the proportion of cells contributed by different major classes. **g,** Pie charts show the proportion of cells contributed by different sub-classes. **h,** Pie charts show the proportion of cells contributed by different donor ancestry. **i,** Pie charts show the proportion of cells contributed by different donor sex.

Supplementary Fig. 3

a

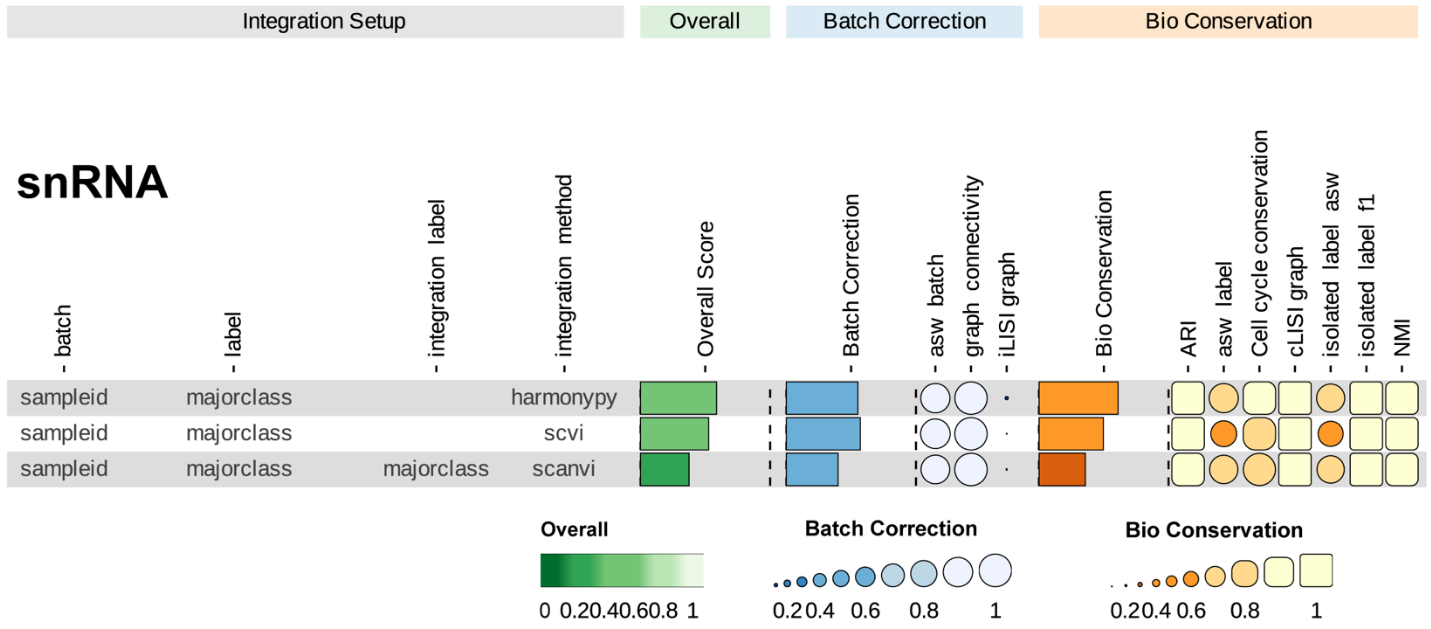


b

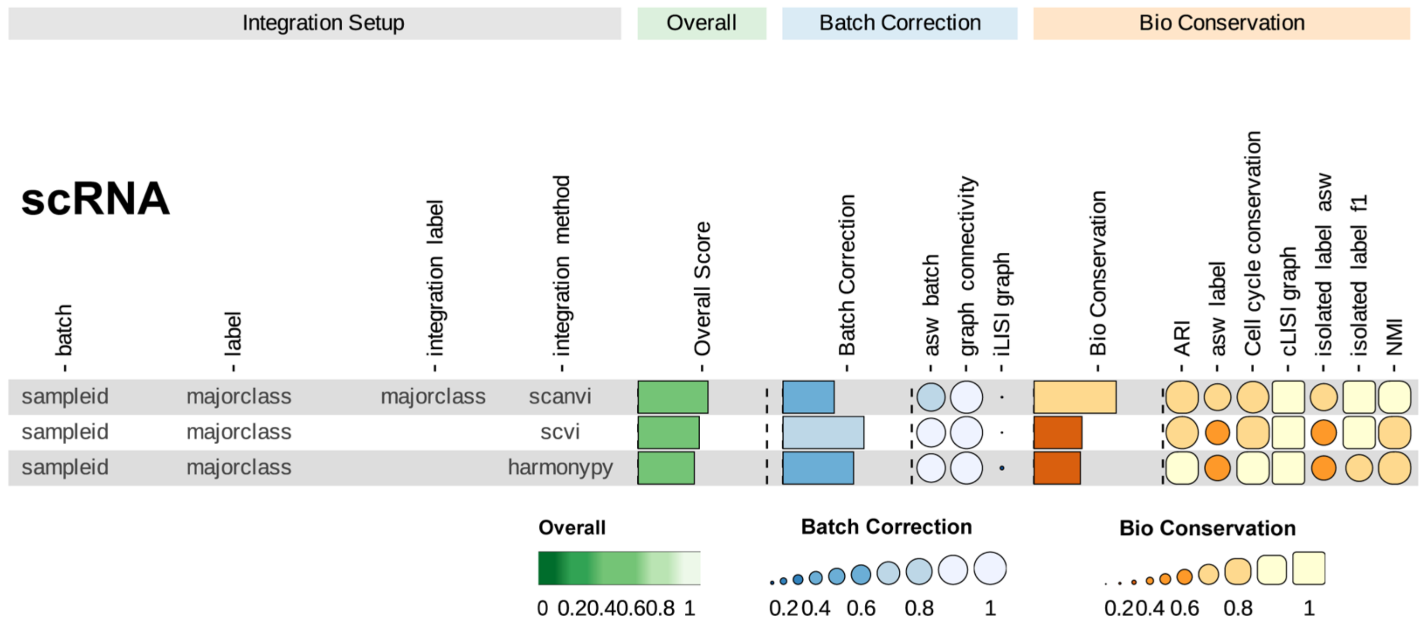


**Supplementary Fig. 3 Benchmark analysis for both scRNA and snRNA atlas of the human TM and CB**

**a,** Benchmark analysis for snRNA atlas. **b,** Benchmark analysis for scRNA atlas.

**Supplementary Fig. 4**

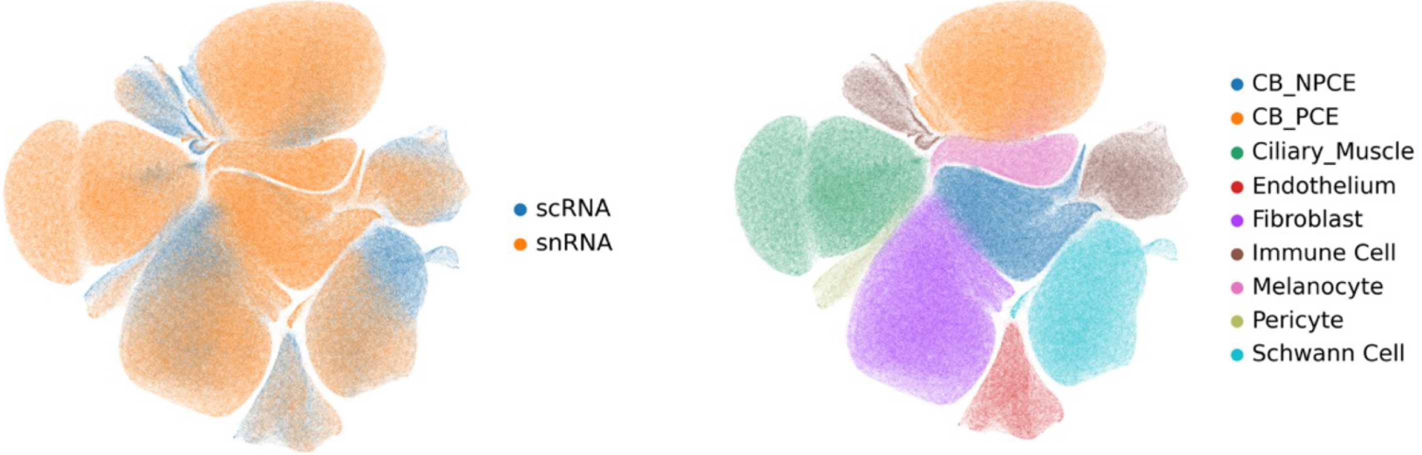


**Supplementary Fig. 4 Integration of scRNA-seq and snRNA-seq datasets**

UMAP visualization of scRNA-seq and snRNA-seq datasets co-embedded using sysVI. Left, cells colored by sequencing modality, showing effective mixing between scRNA-seq and snRNA-seq after integration. Right, cells colored by annotated major classes.

**Supplementary Fig. 5**

**a**

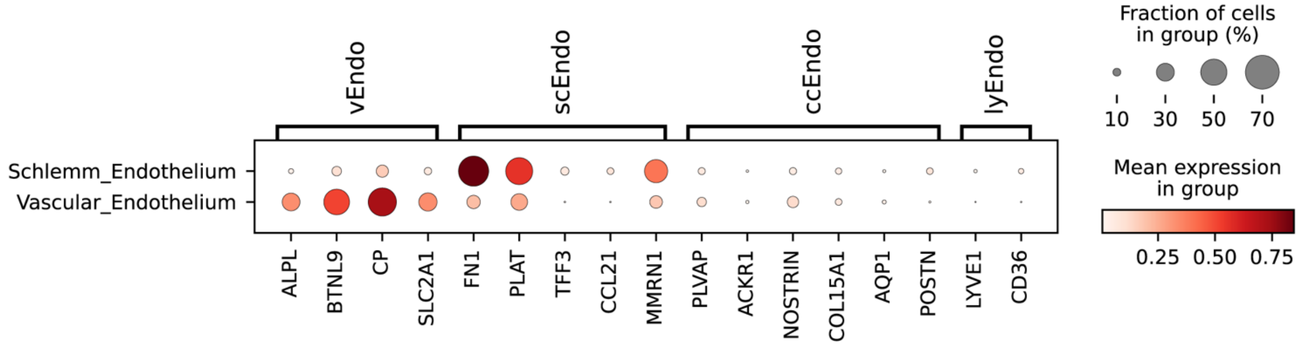


**b**

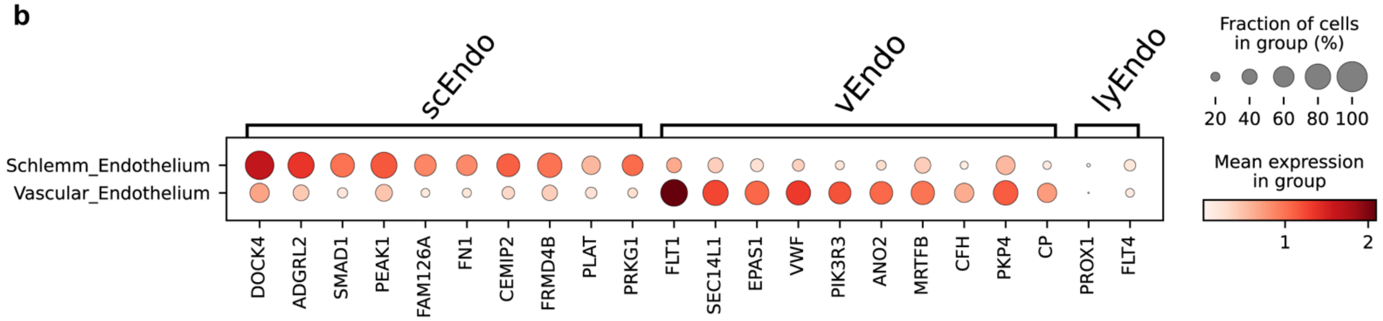


**Supplementary Fig. 5 Dotplot of marker genes expression in Endothelium sub-classes of TM and CB snRNA atlas**

**a,** Known marker genes of different sub-classes expression in Endothelium sub-classes. **b,** Top 10 DEGs expression in Endothelium sub-classes.

**Supplementary Fig. 6**

**a**

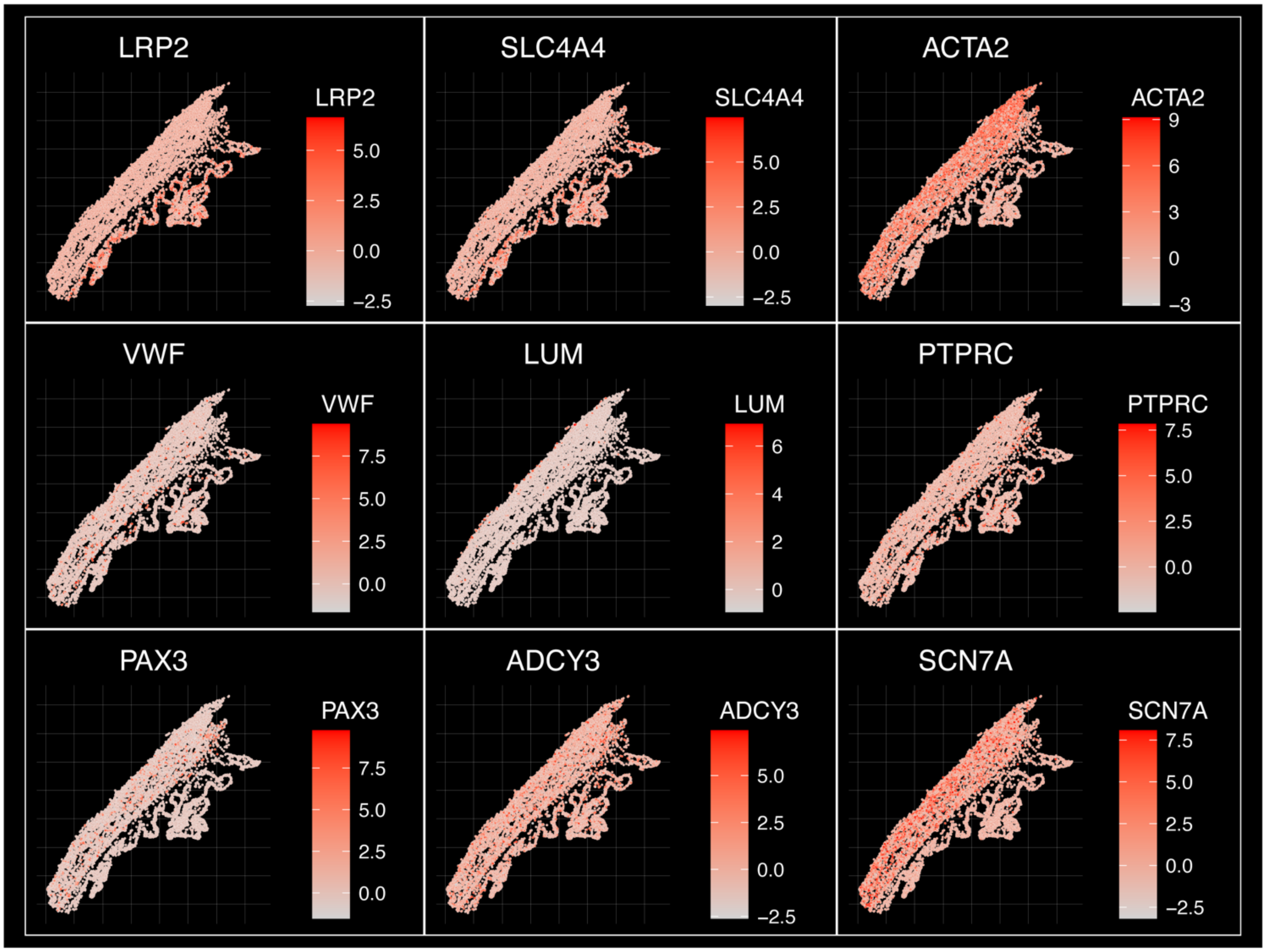


**b**

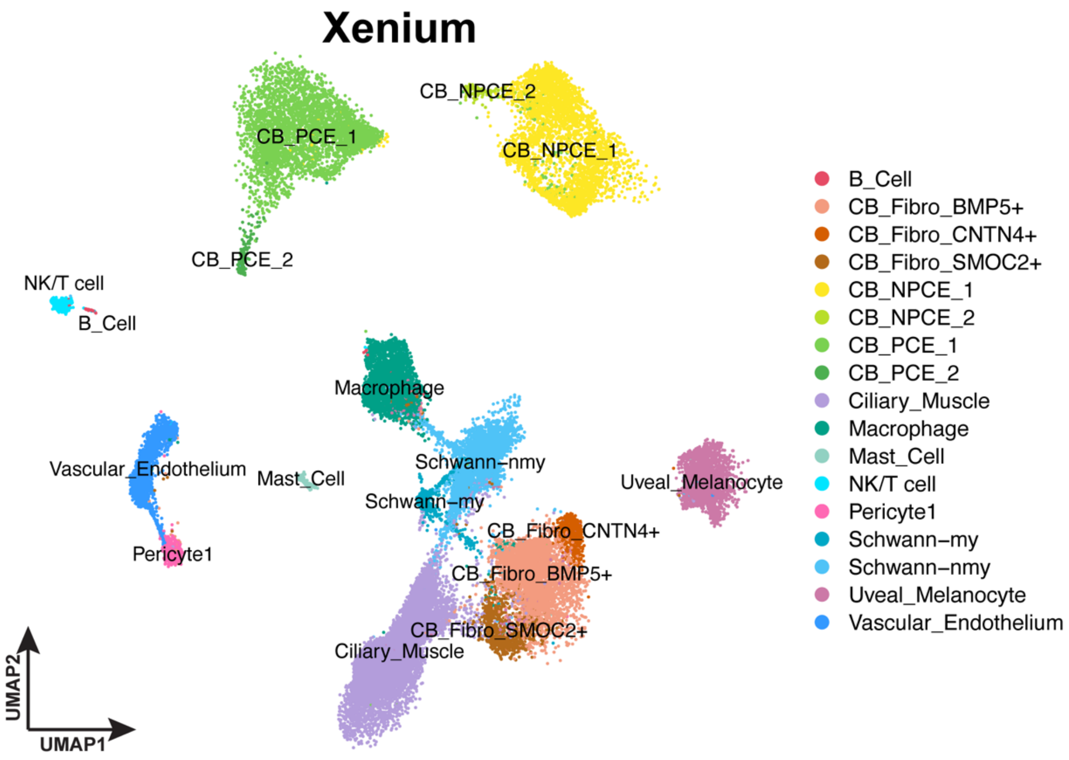

**Supplementary Fig. 6 Marker gene expression and cell type annotation in CB Xenium data**

**a**, Spatial feature plots showing the expression patterns of marker genes across the CB tissue (Slide 1 as example). **b,** UMAP of Xenium data colored by annotated sub-classes.

**Supplementary Fig. 7**

**a**

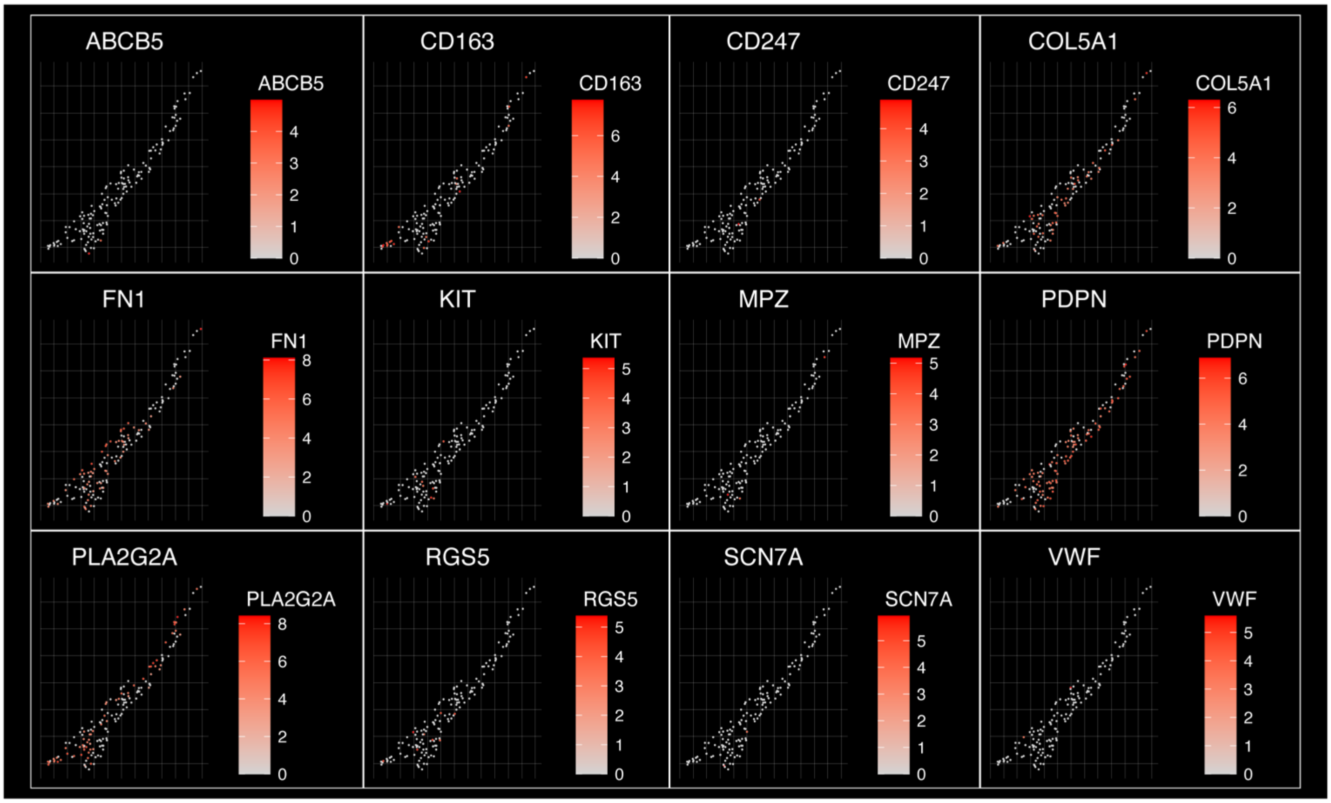


**b**

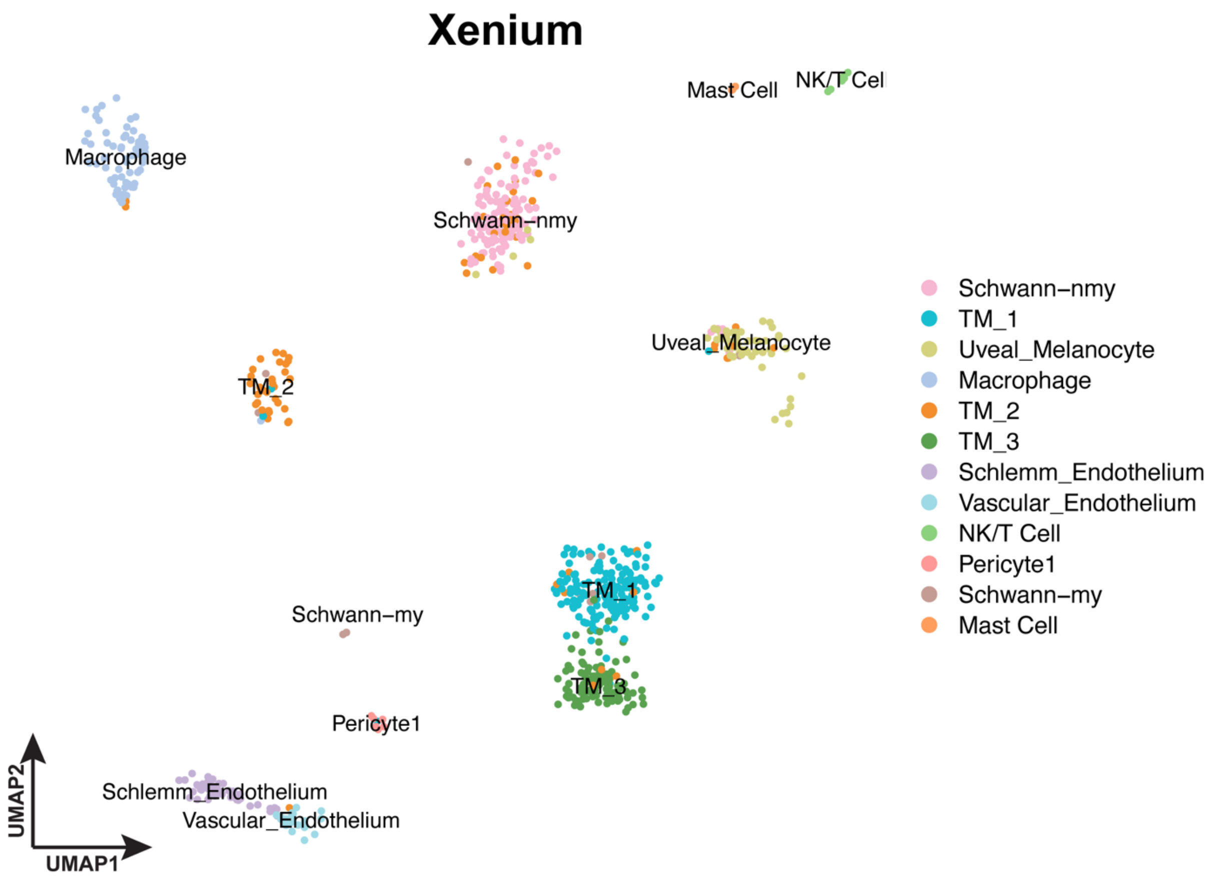

**Supplementary Fig. 7 Marker gene expression and cell type annotation in TM Xenium data**

**a**, Spatial feature plots showing the expression patterns of marker genes across the TM tissue (Slide 1 as example). **b,** UMAP of Xenium data colored by annotated sub-classes.

### Supplementary Fig. 8

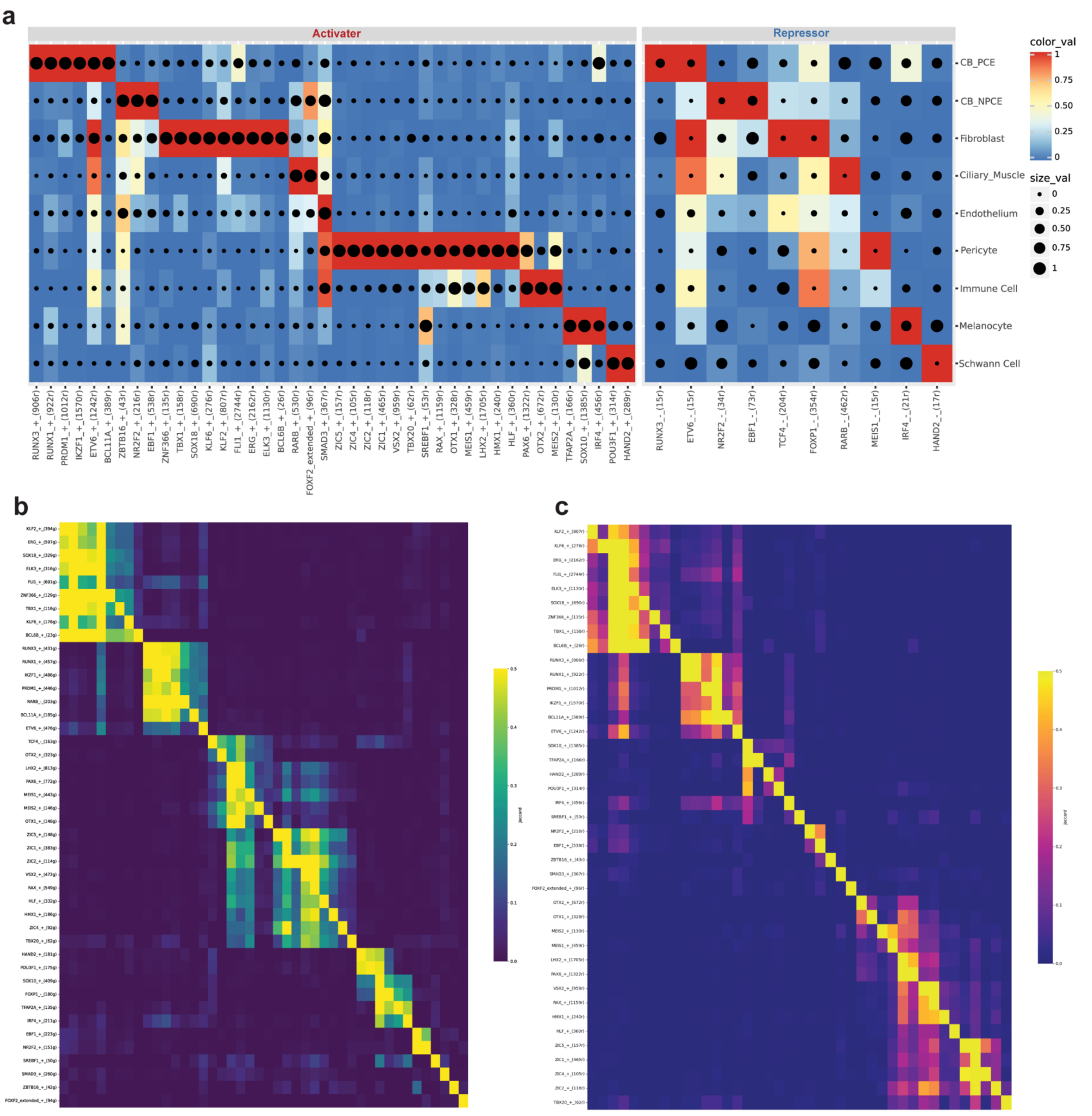

**Supplementary Fig. 8 Transcriptional regulatory programs across cell types**

**a**, Heatmap showing inferred TFs activity across major cell types. Activators (left) and repressors (right) are displayed, with color indicating activity level and dot size representing the proportion of cells with detectable activity. **b–c,** Pairwise correlation heatmaps of transcriptional regulators, showing co-activation patterns and modular organization of regulatory programs across cell populations**.**

### Supplementary Fig. 9

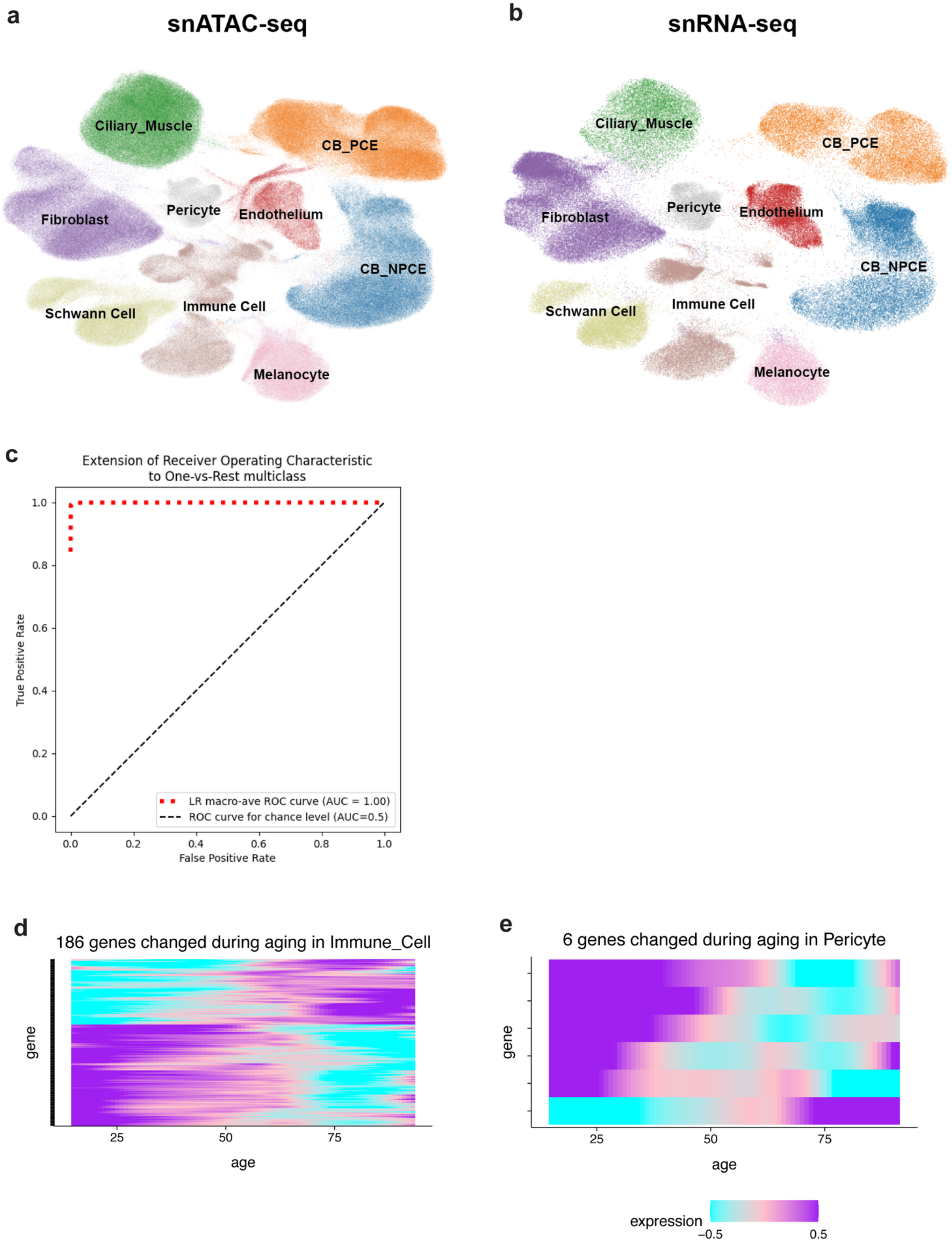

**Supplementary Fig. 9 Transcriptional regulatory programs across cell types**

**a, b,** UMAP of co-embedded cells from snATAC-seq (left) and snRNA-seq (right) showing cells are clustered into sub-cell classes. **c,** Pairwise correlation heatmaps of transcriptional regulators, showing co-activation patterns and modular organization of regulatory programs across cell populations**. d, e,** Heatmap showing gene expression of DEGs during aging in immune Cell (left) and pericyte (right) identified with LMM.

**Supplementary Fig. 10**

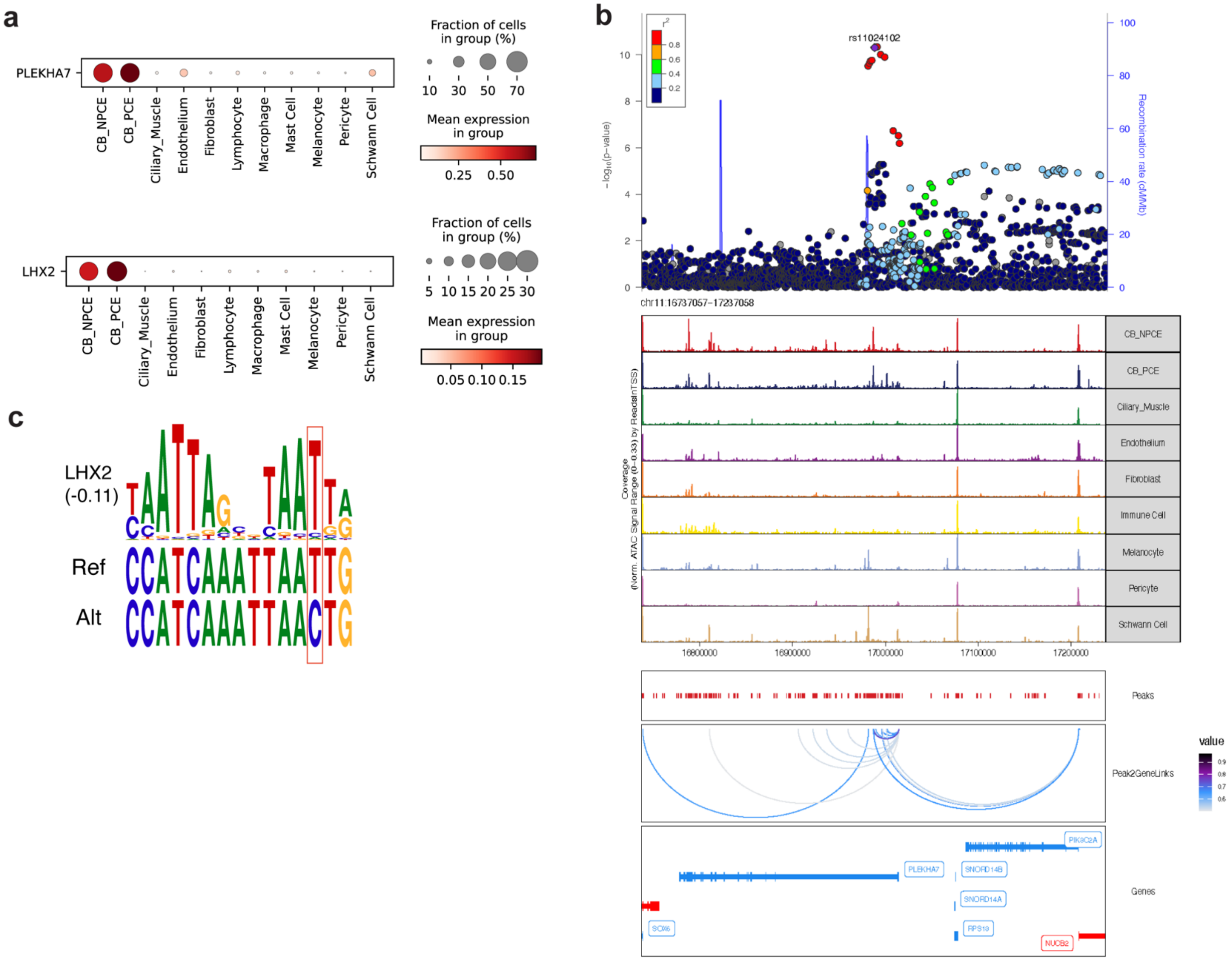


**Supplementary Fig. 10 Fine-mapping and regulatory annotation of the PACG-associated *PLEKHA7*.**

**a,** Expression of *PLEKHA7* and LHX2 across TM and CB majorclasses. **b,** Regional association plot and chromatin accessibility profiles at the *PLEKHA7* locus. The prioritized PACG-associated variant rs11024102 overlaps a regulatory element linked to *PLEKHA7*. **c,** Predicted disruption of an LHX2 binding motif by rs11024102.
